# Live-cell imaging enables reporter-free monitoring of the circadian rhythm in individual *Synechocystis* cells

**DOI:** 10.64898/2026.02.04.703767

**Authors:** Lennart Witting, Florian P. Stirba, Julius Nohr, Ekaterina Ivanova, Petra Kolkhof, Dietrich Kohlheyer, Anika Wiegard, Ilka M. Axmann

## Abstract

In vivo monitoring of circadian rhythms depends on reliable and non-invasive detection methods. This is often achieved by expressing reporter genes heterologously under the control of a circadian promoter. The activity or fluorescence of the gene product is then used as a readout. To avoid the need for generation of such reporter strains, we recently established a reporter-free detection method for cyanobacterial batch cultures. To determine whether these rhythms are driven at the level of individual cells or result from population-based effects, such as gating of cell division, we analyzed individual *Synechocystis* sp. PCC 6803 cells by combining a microfluidic cultivation technique with multipoint time-lapse microscopy imaging at the single-cell resolution. Hundreds of time-lapse image sequences, acquired over a period of up to ten days, were processed using our deep learning cell segmentation workflow. Although the cells had been entrained by a 12-hour light-dark cycle, neither cell size nor cell division displayed circadian rhythms. This indicates the absence of circadian gating of cell division in *Synechocystis*. Instead, we observed circadian oscillation in the average brightness of the phase contrast of individual *Synechocystis* cells. To demonstrate how phase-contrast analysis of single cells can be complemented by backscatter analysis of batch cultures, we investigated the wildtype, a deletion mutant known to affect circadian rhythms (Δ*kaiC3*) and complementation strains at both, the single-cell and batch levels. We concluded that phase contrast and backscatter likely measured the same rhythmic changes in the refractive index of the cells. The method presented here will advance circadian research by enabling the analysis of circadian rhythms in individual cells without the need for expression of reporter molecules.

## Introduction

### Cyanobacterial circadian clocks

Circadian clocks are endogenous timekeeping mechanisms that control biological activities in accordance with the rhythmic day-night regime of the organism’s habitat, which results from the Earth’s rotation. True circadian clocks exhibit temperature compensated oscillation with periods of approximately 24 hours in the absence of external cues [1]. However, the phase of the oscillation can be synchronized with external signals, allowing it to adapt to a rhythmic environment. In cyanobacteria, the circadian clock provides a fitness advantage under rhythmic conditions [2].

In the model organism *Synechococcus elongatus* PCC 7942 (*Synechococcus*), the core oscillator consists of only three proteins: KaiA, KaiB, and KaiC [3, 4]. KaiC is the central component, which drives the circadian cellular responses. It displays 24-hour ATPase activity cycles, as well as rhythmic phosphorylation and dephosphorylation of serine 431 and threonine 432, which are driven by the dynamic interaction with KaiA and KaiB [5–9]. In brief, KaiA binding promotes KaiC autophosphorylation [10, 11] Upon binding to KaiC, KaiB sequesters KaiA from its binding site, thereby facilitating autodephosphorylation of KaiC [12–14]. These three proteins form a precise post-translational oscillator (PTO) that can be reconstituted in vitro and maintains the circadian oscillation in cells without external stimuli and after inhibition of transcription and translation [4, 15]. In *Synechococcus*, a transcription translation feedback loop ensures synchrony within a population without communication between cells [16–19].

Several cyanobacteria and other prokaryotes contain multiple diverged copies of *kai* genes [20, 21]. In *Synechocystis* the clock comprises two KaiABC systems (KaiA1B1C1 and KaiA3B3C3), which together drive circadian rhythms [22]. An additional *kaiB2C2* cluster present in *Synechocystis* does not appear to be involved in circadian regulation in this organism [23]. The KaiA1, KaiB1 and KaiC1 proteins of *Synechocystis* show the highest homology to the respective proteins in *Synechococcus* [20, 21]. KaiA3 was recently identified [22]. It consists of an N-terminal NarL-type response-receiver domain and a C-terminal KaiA domain.

KaiC3 harbors serine 423 and threonine 424 as phosphorylation sites and exhibits dampened phosphorylation cycles in the presence of KaiA3 and KaiB3 in vitro. In vivo, the phosphorylation of KaiC1 and KaiC3 oscillated with the same phase and period. Deletion of either *kaiA3* or *kaiC3* gene, or both, results in a damped oscillation during measurements of cellular functions downstream of the circadian oscillator. Knocking out *kaiB3* or *kaiA1B1C1* completely abolished the oscillation. These results suggested that the dysregulation of KaiC3 phosphorylation significantly impacts the overall circadian oscillation in *Synechocystis* [22]. Even before the discovery of KaiA3, Aoki and Onai suggested that KaiB3 and KaiC3 were involved in finetuning the circadian rhythm of *Synechocystis* [24]. Little is known about how the two PTOs of *Synechocystis* relay information to generate an output.

### Reporter-free detection of circadian rhythms

There are different approaches to measuring circadian oscillations in vivo using non-invasive methods. In *Synechococcus*, the gene-reporter method is frequently applied, which involves the expression of bioluminescence or fluorescence reporter molecules under the control of a circadian promoter so that circadian oscillations in gene expression can be measured. Circadian rhythmicity is monitored by recording bioluminescence or fluorescence over time [18, 25–27]. In such an experiment, cells are entrained by environmental cues (e.g., light-dark rhythms) and gene expression (or other cellular activity) is monitored under free-running conditions (e.g., constant light) [28]. Fluorescence or bioluminescence are measured either for bacterial colonies on plates or for individual cells. The latter can be achieved by culturing cells on agarose pads [18] or in defined microfluidic structures, such as mother machines or 2D cultivation chambers [16, 25–27]. Furthermore, by applying single-cell cultivation methods, researchers have been able to distinguish between cell division and cell elongation. This has allowed them to study how the circadian clock controls cell division in *Synechococcus* [29–31].

In the past, adapting the gene-reporter method to *Synechocystis* has been difficult. This is due to the fact that only 2-7 % of genes appear to be regulated by the circadian clock and the rhythms have a lower amplitude than those in *Synechococcus* [32]. However, Zhao et al. recently detected circadian bioluminescence signals in *Synechocystis* at the colony level using a strong artificial promoter [23]. Other researchers have successfully cultivated *Synechocystis* in PDMS-based microfluidic photobioreactors and recorded its growth at the single-cell level. However, to the best of our knowledge, these publications did not include observations of circadian rhythms [26, 33].

An alternative to the gene-reporter method is monitoring circadian rhythms in batch cultures using non-invasive, online backscatter measurements. This method does not require genetic manipulation or knowledge of a suitable promoter for expressing a reporter molecule and is therefore referred to as ‘reporter-free’. This is advantageous because in *Synechocystis* it is unclear which PTO output components (e.g. RpaA, SasA and CikA) control which promoter and how these are connected to the two KaiABC systems. We previously reported such oscillations in *Synechocystis* and *Synechococcus* wildtype batch cultures and we demonstrated that oscillations meet all three criteria of circadian rhythms. Notably, synchronization of the batch culture could also be achieved by dilution and did not require light-dark entrainment [34]. However, it is not yet known whether the backscatter oscillation detects changes in the optical properties of individual cells or batch-culture effects, such as circadian cell division or cell elongation.

In this study, we combined the advantages of the backscatter method with those of the single-cell cultivation methods established for *Synechococcus*. We used single-cell cultivation technology to trap and image individual cells over multiple days to gain insights into the circadian clock output of *Synechocystis* at the single-cell level. Using our cell segmentation workflow, we analyzed single-cell resolution time-lapse image data and revealed circadian rhythms in the phase-contrast intensities of individual cells. The single cell oscillations exhibited the same phase and period across the population. No evidence of cell division gating was found, and the colonies grew exponentially without a superimposed oscillation. As with the backscatter oscillation [22], knocking out the *kaiC3* gene resulted in a damped phase-contrast oscillation. Additionally, using complementation strains, we demonstrated that amino acid exchanges in KaiC3 can manipulate the backscatter oscillation. Because both methods of detecting the circadian rhythm are influenced by the refractive index of individual *Synechocystis* cells, we conclude that they must measure the same phenomenon on different population scales.

## Results

### Single-cell cultivation enables reporter-free observation of the circadian rhythm in individual *Synechocystis* cells

Although the oscillation of the backscatter in batch cultivation is linked to the circadian clock, identifying the phenomenon causing the oscillation has been difficult using the backscatter technique. Crucially, because backscatter measures a cumulative signal from a large cell population, it was not possible to determine whether the oscillation was caused by population-level phenomena (e.g., timing of cell division) or by changes occurring within each cyanobacterial cell simultaneously (e.g., changes in the refraction properties of individual cells). To further investigate this, we used single-cell cultivation and time-lapse imaging with single-cell resolution, which enabled us to separate and observe individual cyanobacteria for up to ten days. Furthermore, single-cell imaging allowed for the observation of cell growth and changes in cell phenotype simultaneously, thus dissecting both phenomena.

*Synechocystis* wildtype (WT) cells were entrained in batch culture under 12 h light followed by 12 h darkness. After the end of the 12 h night, the cells were transferred into the growth chambers on a microfluidic cultivation chip. The time required to inoculate the cells, select suitable positions for time-lapse imaging, and initiate the time-lapse experiment varied from one experiment to the next, but never exceeded two hours. The cultivation chip was illuminated with constant white light (Fig. S1) at 4 µmol_(photons)_·m^-2^·s^-1^ intensity and the cells were imaged every hour (Fig. 1A).

**Figure 1:**
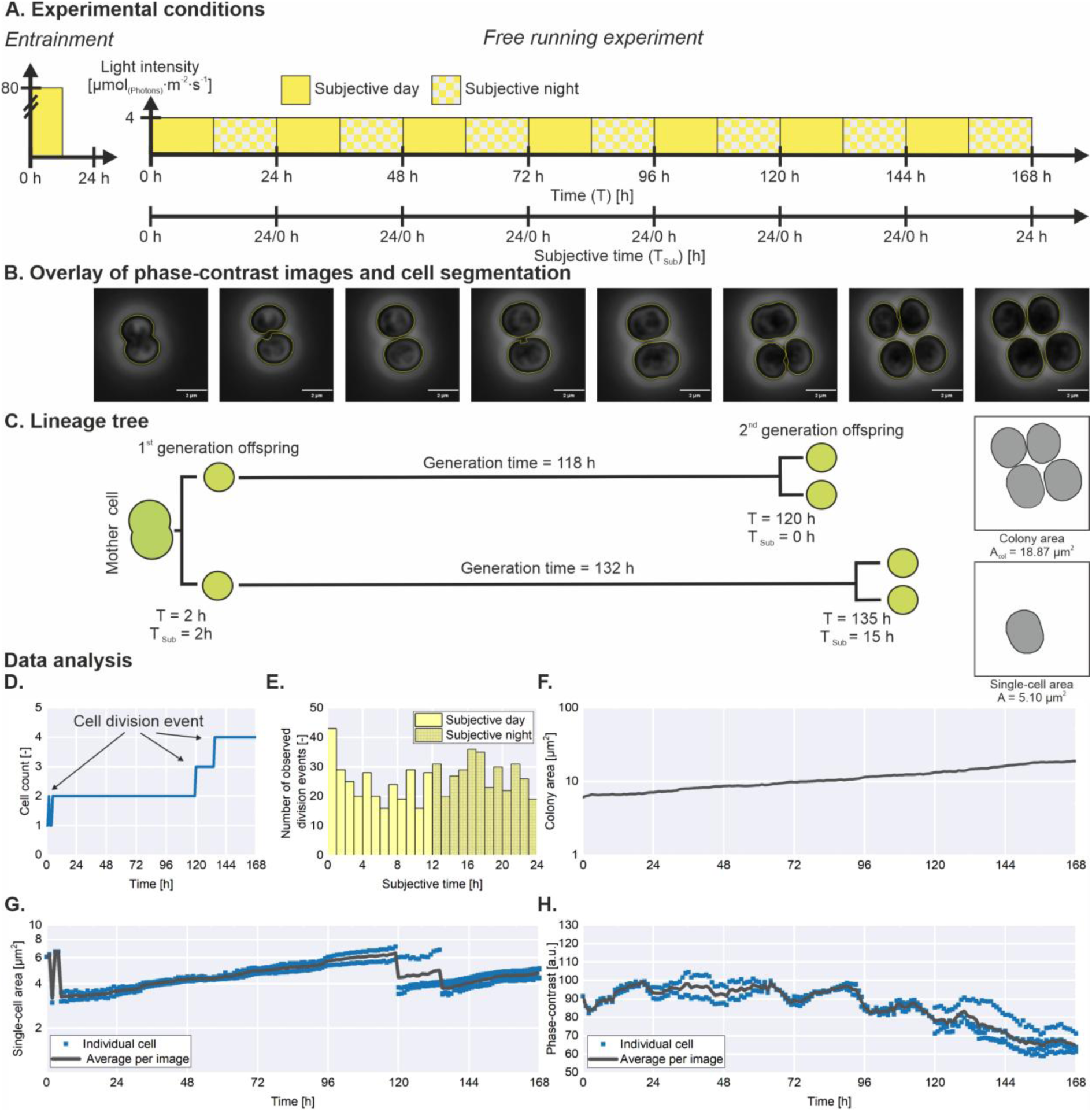
Microfluidic, single-cell-level cultivation enables the observation of circadian rhythms. Results from an exemplary WT colony are shown. A: The strains were precultivated and synchronized in a photobioreactor. During single-cell cultivation in a microfluidic chamber, constant white light at 4 µmol(photons)·m^-2^·s^-1^ was applied. The 168 h of cultivation time (T) is subdivided into seven subjective days (TSub). Each subjective day consists of a 12-h subjective day and a 12-h subjective night. B: During time-lapse microscopy images were captured every hour. Here, example images (24 interval) of the *Synechocystis* WT growing in a microfluidic cultivation chamber are displayed. Cells were segmented using a previously published data-processing pipeline [35]. C: Lineage trees were reconstructed by visually inspecting the image sequences. D: Plot of the number of cells in an image (cell count) over time. E: These plots can be used to analyze the subjective time at which cell division occurred. A total of 624 division events were detected in 141 image sequences from nine independent experiments and plotted over the subjective time. F + G: Furthermore, the method allows for the distinction between the size of individual cells (single-cell area) and the cumulative area occupied by a cell colony (colony area) H: Additionally, the brightness of individual cells within the phase-contrast images (phase-contrast) can be extracted.

To quantify changes within single cells over time, the *Synechocystis* cells within time-lapse images were segmented using our previously published cell segmentation workflow [35] (Fig. 1B).

Due to the small number of cells, lineage trees (Fig. 1C) could be easily reconstructed by manually inspecting time-lapse videos. The time point at which cell division occurred was automatically extracted from the plot of the number of cells in a frame of an image sequence (cell count) over time (Fig. 1D). Data from 141 cell-count plots from nine independent single-cell experiments, equivalent to the one shown in Fig. 1D, were investigated. A total of 624 division events were identified. This revealed that cell divisions occurred equally distributed over the whole 24h day under free-running conditions (constant light) (Fig. 1E), suggesting that cell division was not timed by the circadian oscillator.

In addition to observing cell division events, the segmentation workflow allows the extraction of time-resolved data on the projected cell surface area of individual cells (single-cell area) and the cumulative area of a colony (colony area), from which the growth rates were derived. The increase in cell area was exponential and not superimposed by an oscillation. This was true for individual cells (Fig. 1G) as well as for the cumulative colony area (Fig. 1F).

However, when we analyzed the average brightness of all pixels composing individual cells (phase-contrast), we observed an oscillation with a period of ∼24 h (Fig. 1H). Notably, this oscillation was observed at the level of individual cells (blue dots) and in the averaged data for all cells within an image (black line), demonstrating robust oscillations in individual cells. The timepoints at which cell division occurred (Fig. 1C) did not correspond to the maxima and minima in the phase-contrast plot. This excludes the cell division cycle as a cause of the oscillation in cell brightness. Fig. 2 shows similar plots for additional image sequences acquired during the same experiment. These plots demonstrate that the phase-contrast oscillation was present in all observed WT cells and that the phase of the oscillation was consistent between colonies and between mother and daughter cells. These findings imply that the observed oscillation during batch cultivation might be caused by oscillations in individual cells rather than population-level effects.

**Figure 2:**
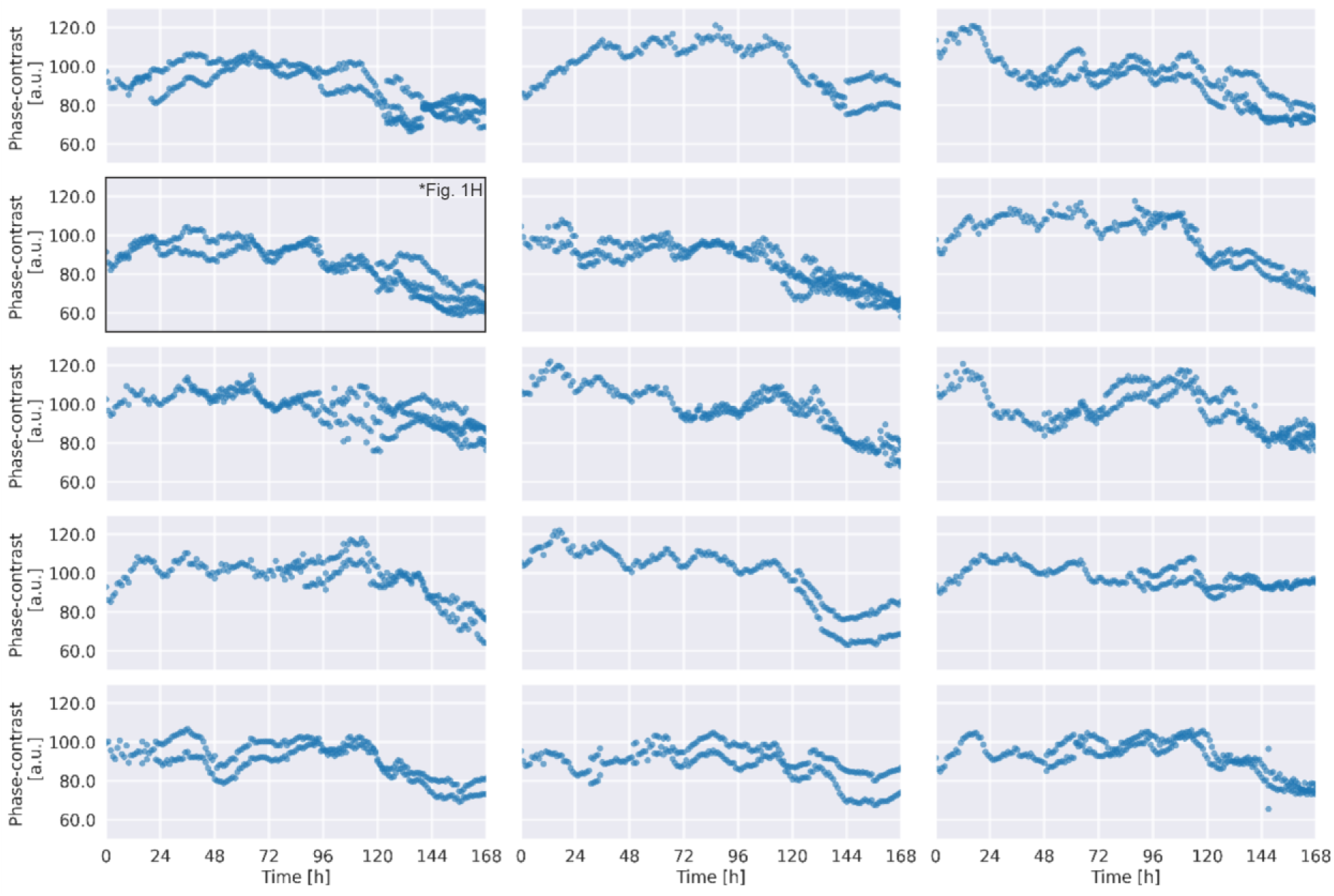
Single-cell phase-contrast intensity from 15 image sequences of the WT. Each plot contains data from a separate image sequence of a single colony acquired during a continuous microfluidic experiment. Although each colony was located in a separate growth chamber, all growth chambers belonged to the same array of chambers on the microfluidic chip. Therefore, they are interconnected by shared supply channels. Each data point represents the mean intensity of all pixels within the boundary of an individual *Synechocystis* cell. The plot in the second row of the first column was previously shown in Fig. 1H.

### A data analysis pipeline for comparing circadian rhythms observed using batch and single-cell cultivation

To test the hypothesis that the population-level oscillation represents single cell oscillations, we investigated whether the phase shifts and periods observed in individual cells were similar to the population-level oscillation. Alternatively, the signal observed in individual cells may be masked when the average oscillation of all cells observed during single-cell cultivation is calculated. Therefore, the single-cell phase-contrast intensities were averaged per frame (Fig. 3A) to show the mean oscillation of all daughter cells within a colony. A data analysis pipeline based on previous publications [22, 34] was designed to further analyze the signals. To isolate the oscillation, a polynomial regression was fitted to the average signal (Fig. 3A), and the relative phase-contrast was calculated by subtracting the raw data from the polynomial fit. After smoothing the data using a sliding window average, robust oscillation within a colony was confirmed (Fig. 3B). The oscillation remained apparent when the mean relative phase-contrast of all image sequences was calculated (Fig. 3D). This confirmed that all the individual cells oscillated with approximately the same phase throughout the experiment. Discrete Fourier transformation (DFT) was performed on the mean relative phase-contrast from 15 independent colonies, and a dominant period of 24 h with an amplitude of 1.56 a.u. (Fig. 3C) was determined for this example. Then a cosine curve was fitted to the mean signal using these values as an initial guess. The resulting amplitude, frequency and phase of the optimized cosine wave were used for further analysis.

**Figure 3:**
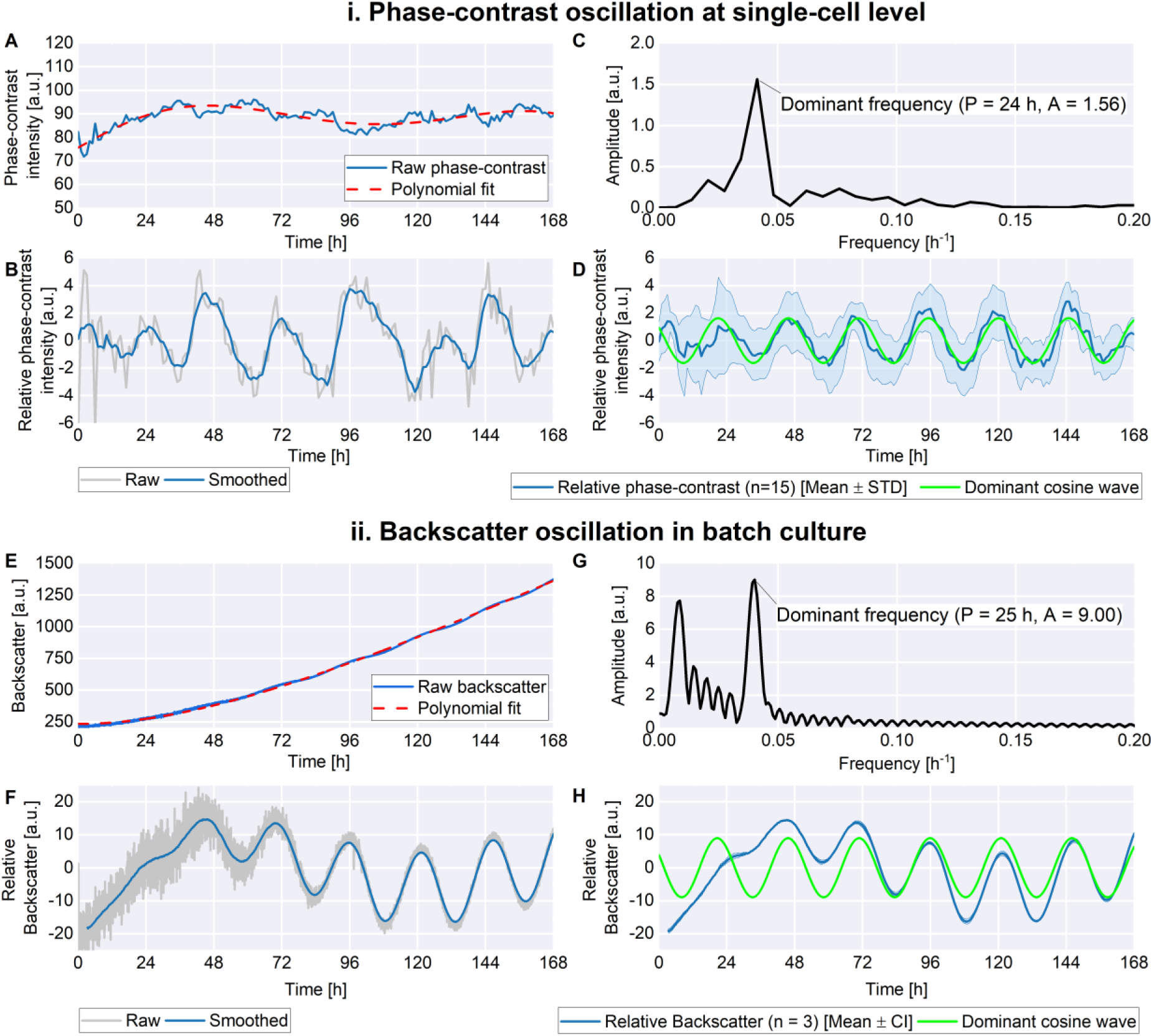
The circadian data analysis pipeline for (i) averaged single-cell and (ii) backscatter data on the *Synechocystis* WT. A: Raw phase-contrast intensity (blue) and polynomial fit (red) of an example image sequence (different example than Fig. 1H). B: The relative phase-contrast (gray) was obtained by subtracting the raw phase-contrast intensity from the polynomial fit (Fit – raw data). A moving average filter for data smoothing was then applied. C: Discrete Fourier transformation performed on the mean relative phase contrast of 15 chambers. The frequency with the highest amplitude is the dominant frequency. D: Mean Relative phase contrast of 15 chambers (blue) and standard deviation (light blue) as well as the reconstructed dominant cosine wave (green). E: Raw backscatter signal (blue) and polynomial fit (red) of an example batch cultivation experiment. F: The raw relative backscatter (gray) was obtained by subtracting the polynomial fit from the raw backscatter (Raw data - fit). Note, that the relative phase-contrast (A-D) was calculated in the opposite way (Data – fit), This was done to ensure that the relative maxima and minima align for both methods, as raw backscatter and raw positive phase-contrast are inversely correlated. Next, a moving average filter was applied to smooth the relative data (blue). G. Discrete Fourier transformation performed on the smoothed data. H: Mean relative backscatter of three cultivations (blue) and the reconstructed dominant cosine wave.

To compare the single-cell measurements with the oscillation in batch culture, we performed backscatter experiments in parallel to single-cell experiments, using cells from the same agar plates for both methods. The cultures inoculated from the plates were incubated for 7 days, diluted once and incubated for further 3 days. After a final dilution, the backscatter measurement was started. To process the raw backscatter data, a similar pipeline as for the microfluidics measurements was constructed to keep the results of both methods comparable. For each replicate, a polynomial regression was fitted to the raw backscatter data (Fig. 3E). The residuals were calculated, normalized by subtracting the arithmetic mean, and smoothed with a sliding window average (Fig. 3F). On the mean of the resulting smoothed signals (n=3) DFT (Fig. 3G) was performed (T=25 h; A=9.0) and with the resulting oscillation parameters a cosine oscillation was fitted to the average smoothed signal (Fig. 3H). Again, the amplitude, frequency and phase of the optimized oscillation were used for further analysis.

During single-cell cultivation the WT exhibited a period of 24.99 ± 0.65 h, an amplitude of 1.35 ± 0.43 and a phase of 3.81 ± 1.97 h (average ± SD of 9 experiments). The time at which the first peak of the cosine oscillation occurred was calculated from these values according to Equ. 10. The calculation showed that the first peak occurred, on average, 21.19 ± 1.54 h after the experiment began. Thus, it occurred towards the end of the subjective night. For the backscatter a period of 25.1 ± 0.02 h, an amplitude of 8.93 ± 0.54 and a phase of 4.51 ± 0.16 h (average ± SD of 3 replicates) was measured. The first peak occurred 20.59 ± 0.14 h after starting the backscatter measurement. While the phase of the cosine wave is dependent on experimental factors (like the time between inoculation of the cultures and the start of the experiment), the period is independent of that. Therefore, it is remarkable that the periods of the WT in both experimental setups were quite similar. As the amplitude is dependent on multiple factors such as sensitivity of the measurement or gain settings, it is not a suitable parameter to compare between different experimental setups. Altogether the similarity of phase and period measured in the two different experimental set-ups support the hypothesis that the individual oscillations are similar enough to cause the previously observed oscillation in backscatter during batch cultivation.

It is worth mentioning that the respective relative signals were calculated in the opposite way. The amount of scattered light per cell is a function of the cell’s refractive index [36–38]. The refractive index can be correlated with the volumetric density of a probe and subsequently with the optical properties of the bacteria [37]. The higher the refractive index, the higher the backscattered light intensity. Lower positive phase-contrast intensity (dark areas) indicates a higher refractive index and volumetric density of the sample. Accordingly, the positive phase-contrast intensity and backscatter are inversely correlated.

### Application of the data analysis pipeline (i) Single-cell cultivation

To prove that the 24h single-cell oscillations were driven by the circadian clock and not artifacts caused by unknown environmental changes in the laboratory (e.g. humidity or temperature), we investigated a *Synechocystis kaiC3* knockout strain (Δ*kaiC3*). This is because the Δ*kaiC3* deletion mutant exhibits a dampened oscillation with decreasing amplitude in batch culture [22]. Accordingly, *kaiC3* is not only an essential component of the *Synechocystis* clock, but it’s deletion also provides a suitable way to estimate the sensitivity of oscillation analysis.

Clearly, a damped oscillation resulting from knocking out *kaiC3* was also observed in the phase-contrast (Fig. 4B). For this strain the period was inconsistent between the representative experiment shown in Fig. 4B (21.01 h) and the replicate experiment shown in Fig. S2 (64.4 h). However, since the data analysis pipeline was optimized to detect a simple harmonic oscillation, not a damped oscillation, this was likely an artifact. To exclude any secondary effects in the *ΔkaiC3* strain, we generated a complementation mutant, in which the wild-typical *kaiC3* gene was reintroduced in the native chromosomal location and expressed under the *kaiC3* promoter. For selection, a spectinomycin resistance cassette was inserted downstream of *kaiC3*, which could affect transcript and protein levels. However, using a KaiC3 specific antibody we detected similar KaiC3 protein levels as in the WT (Fig. S11). Analysis in the microfluidic setup confirmed that re-insertion of the *kaiC3* gene restored the single-cell oscillations (Fig. 4C). The periods (26.29 and 25.80 h), amplitudes (2.18 and 2.11 h), phases (4.46 and 4.48 h) and first peaks (21.83 and 21.42 h) were similar in two experiments analyzing the *kaiC3-ST* complementation strain (Fig. 4C and Fig. S2), and were consistent to the values observed for the WT. To investigate the oscillation behavior of the Δ*kaiC3* strain further, we examined the single-cell oscillation plots of the Δ*kaiC3* and the *kaiC3-ST* strain (Fig. S3 and S4). Additionally, we calculated the oscillation parameters of all strains for each image sequence individually, instead of for the mean from multiple image-sequences (Fig. S5). The data in these Figures demonstrate that knocking out *kaiC3* led to dissimilar single-cell oscillation phenotypes within populations. Although some *ΔkaiC3* cells exhibited phase-contrast oscillations, the amplitudes were lower than those observed in the WT and the oscillations were inconsistent within a colony. Other cells did not exhibit oscillations at all. Overall, this variation in phenotype results in the apparent damped phenotype, as seen in the averaged colony data. Similar to the WT, no superimposed circadian oscillation was found in the plots of colony area over time for the knockout and complementation strain (Fig. S6). As shown in Fig. 4E, the area of a single cell was more than twice the height of the cultivation chamber (1.72 µm) during single-cell cultivation. Thus confirming, that the *Synechocystis* cells were grown in a strict monolayer. Therefore, the increase in the area that a colony occupies must be directly proportional to the cell expansion. The growth rates were slightly lower for the WT than for the knockout and complementation mutants. The calculated rates correspond to mean doubling times of 106.99 ± 20.14 h (n = 15), 91.72 ± 14.15 h (n = 14), and 84.98 ± 7.00 h (n = 14) for the WT, knockout, and complementation mutants, respectively. The cause of the slight deviation in growth rate remains unclear; however, it is evident that the cell division cycle is not responsible for the oscillation. Furthermore, the knockout did not affect the area distribution of the single cells. This is noteworthy because if the knockout reduced cell size, it could affect the microscope’s ability to detect changes in cell brightness. As an additional control, we cultivated the WT illuminated with 7 µmol_(photons)_·m^-2^·s^-1^ instead of 4 µmol_(photons)_·m^-2^·s^-1^. The growth rate was higher at 7 µmol_(photons)_·m^-2^·s^-1^ (0.011 ± 0.002 h^-1^) than at 4 µmol_(photons)_·m^-2^·s^-1^ (0.007 ± 0.001 h^-1^). However, the difference in oscillation parameters did not exceed differences observed between independent experiments at the same light intensity (Fig. S7). Hence, after analyzing the single-cell data acquired for the WT, the *ΔkaiC3* and the *kaiC3-ST* strain, it is evident that the oscillations observed in individual cells are linked to the circadian oscillator.

**Figure 4:**
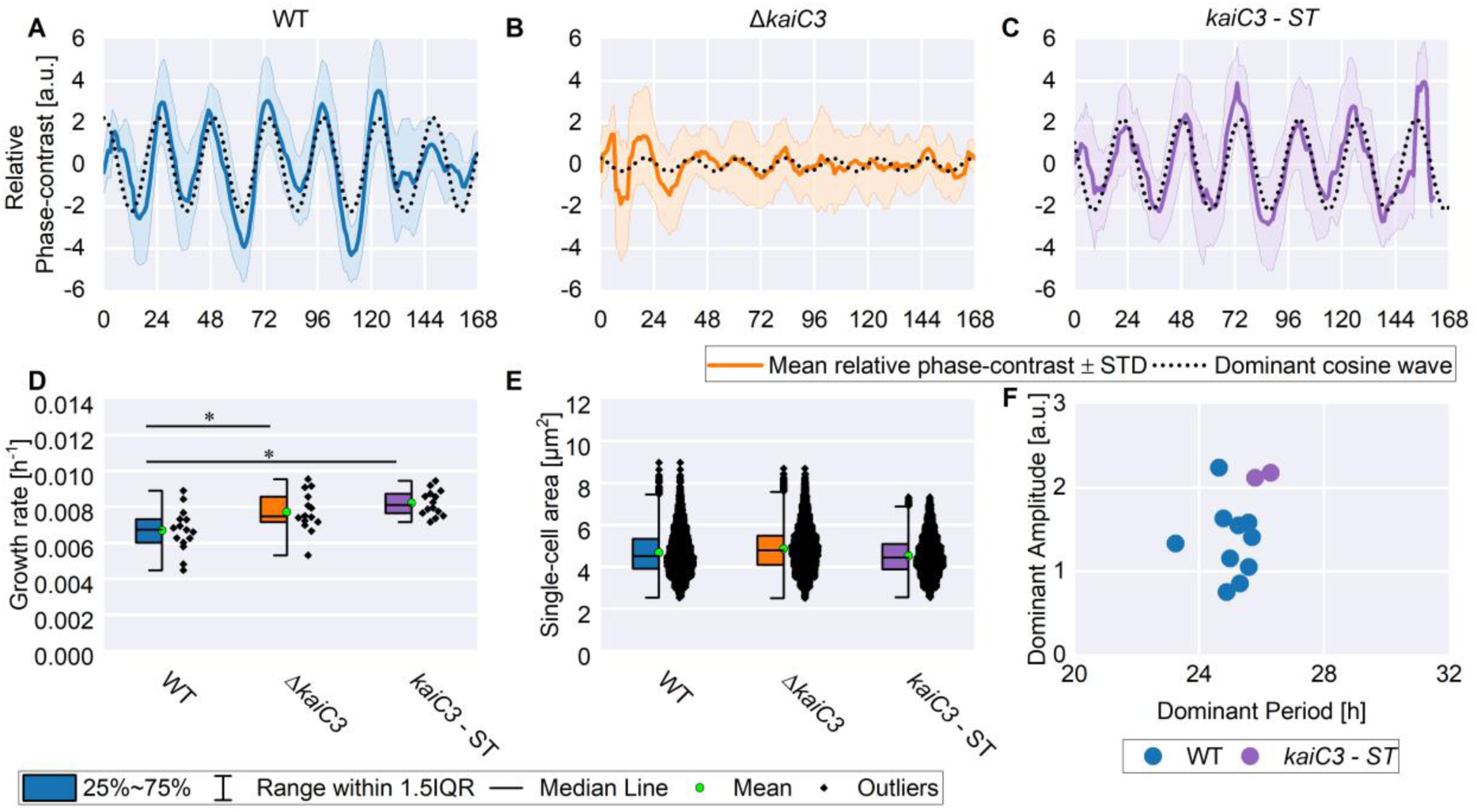
Analysis of the circadian oscillation of the phase-contrast during single-cell cultivation. A-C: Mean relative phase-contrast intensity and the dominant cosine wave for the WT (blue; average of the data shown in Fig. 2), Δ*kaiC3* knockout (orange) and the *kaiC3-ST* complementation mutant (purple). D: Boxplots displaying the growth rates derived from colony area plots E: Distribution of the single-cell areas during single-cell cultivation for the WT, the Δ*kaiC3* knockout and the *kaiC3–ST* complementation mutant. The growth rates were compared by one-way ANOVA followed by Tukey’s HSD test (* p < 0.05). F: Period and amplitude of the optimized dominant cosine wave. In F each data point represents growth within a distinct growth array on a microfluidic chip. A total of nine experiments were conducted with the WT. In one of the nine experiments two arrays were inoculated with the WT, therefore ten data points are shown for the WT. For both mutants, a repeat experiment was performed (Fig. S2).

### Application of the data analysis pipeline (ii) Batch cultivation

As demonstrated in the previous sections, individual WT cells oscillated with periods and phases comparable to those observed in backscatter measurements and *kaiC3* deletion dampened the oscillation in individual cells. To compare oscillation parameters between single cells and batch cultures, we also investigated Δ*kaiC3* and *kaiC3-ST* using the backscatter method. Fig. 5 shows the results of one representative backscatter experiment. The isolated backscatter signal represents the arithmetic mean of 3 replicates with the 95 % confidence interval (CI) shown as error. Data from each triplicate are shown in the Supplement (Fig. S9-S10). The period (frequency), phase and amplitude of the oscillation was determined by DFT for each replicate separately. The results from this analysis were used as initial parameters to fit a cosine curve to the isolated oscillation signal (Fig. 5A – D). From the cosine fit, the values for period, amplitude and phase displayed in Fig. 5 E and F were determined. The WT (Fig. 3H) and the *kaiC3-ST* (Fig. 5B) strain exhibited a circadian oscillation with similar periods (WT: 25.09 ± 0.02 h; *kaiC3-ST*: 25.40 ± 0.01 h), which were also similar to the periods detected in single cells. Except for the phase, WT-like oscillation was restored in the *kaiC3*-ST complementation strain. when analyzed in batch culture. The average phase of the complementation strain was lower (WT: 4.51 ± 0.16 h; *kaiC3-ST*: 1.05 ± 0.04 h), while the amplitude increased (WT: 8.93 ± 0.44; *kaiC3-ST*: 12.95 ± 0.12). The first oscillation peak appeared 20.59 ± 0.14 and 24.34 ± 0.03 h after the start of the measurement for the WT and the *kaiC-ST* complementation mutant, respectively.

**Figure 5:**
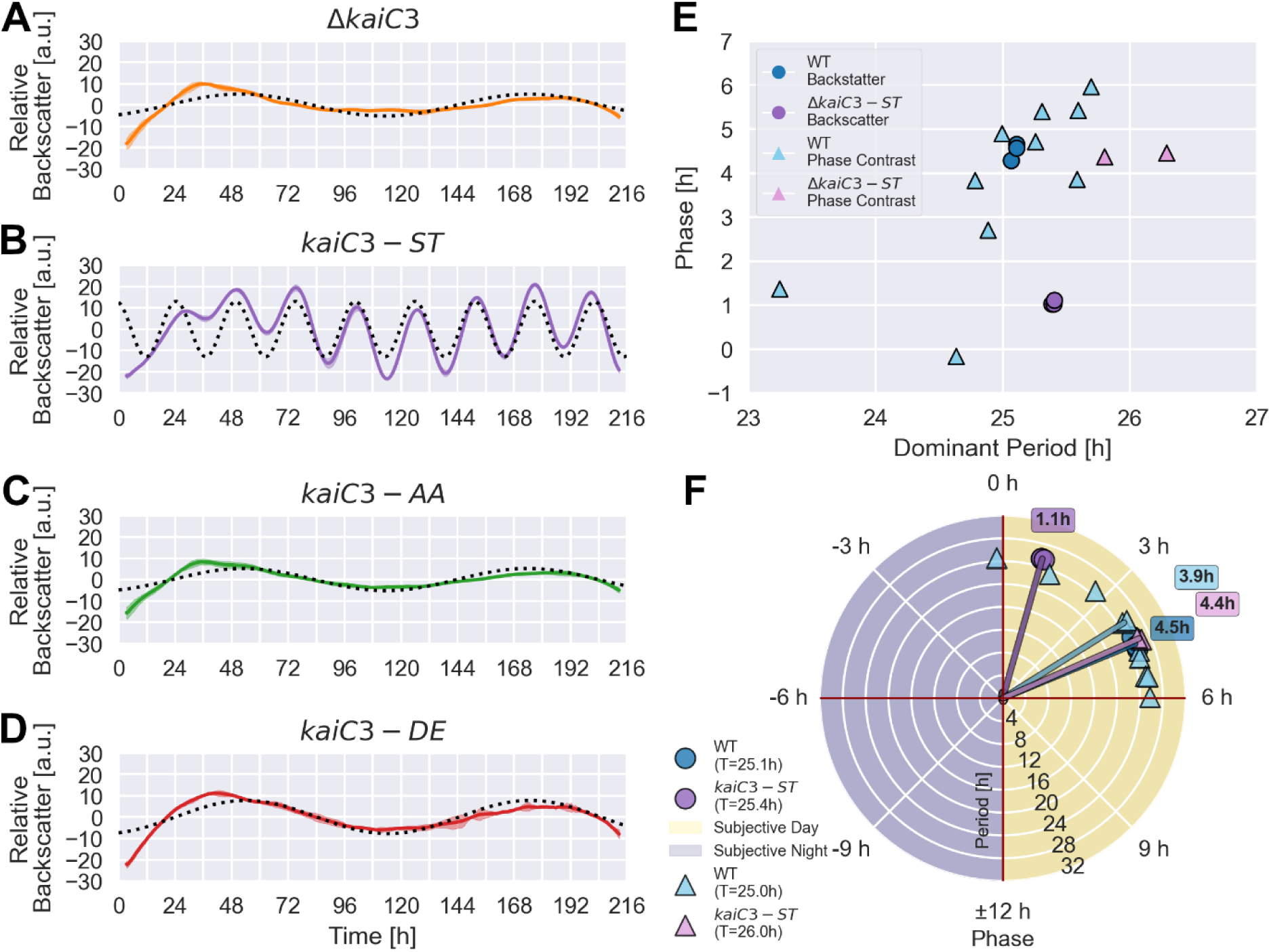
Analysis of the circadian oscillation of the backscatter during batch cultivation. A – D: Mean relative backscatter measurements (n = 3; solid lines) of the *kaiC3* knockout (Δ*kaiC3*), the *kaiC3* complementation strain (*kaiC3-ST*) and the *kaiC3* phosphomimetic mutants (*kaiC3-AA* and *kaiC3-DE*). The 95 % confidence interval (CI) based on a t-distribution was used as an indicator for the uncertainty of the measurement (shade). To each mean backscatter curve, a cosine function (dotted line) was fitted using the amplitude, period (frequency) and phase determined by discrete Fourier transformation as initial guesses for these parameters. E: Scatter plot comparing the dominant period and phase derived from batch culture (backscatter) and single cell (phase contrast) analysis. F: Polar plot of the mean phase (phase vector) and the individual replicates for the WT and the complementation mutant. The radius indicates the period of the oscillation. In E and F the backscatter (Circle) oscillation parameters were derived for all three batch culture replicates considered for Fig. 3H (WT) and Fig 5B (*kaiC3*-ST) individually. For the phase contrast (triangle) each datapoint represents growth within a distinct growth array during a single-cell experiment.

To demonstrate that the oscillation is governed by the phosphorylation status of KaiC3, rather than by the abundance of the protein itself, we further investigated two phosphomimetic mutants (*kaiC3-AA* and *kaiC3-DE*), where the amino acid residues of KaiC3 that can be phosphorylated (S423 and T424) were exchanged by AA (simulating constant dephosphorylation) or DE (simulating constant phosphorylation) (Fig. 6 and 5C - 5D). In both phosphomimetic mutants circadian rhythms were abolished like in the Δ*kaiC3* mutant, proving that the Serine 423 and Threonine 424 are essential for the circadian output of the whole cell even though the KaiA3B3C3 oscillator is only partly responsible for keeping the rhythm [22]. These findings connect the changed genotype (exchange of two amino acids) with a measurable phenotype of the backscatter properties (cessation of oscillation). Furthermore, the experiment outlines how our two methods can complement each other. Single-cell cultivation allows for detailed observation of the oscillation in individual cells; however, only a few strains can be compared. The batch cultivation method, on the other hand, allows us to screen and compare the oscillation parameters of multiple mutant strains.

**Figure 6:**
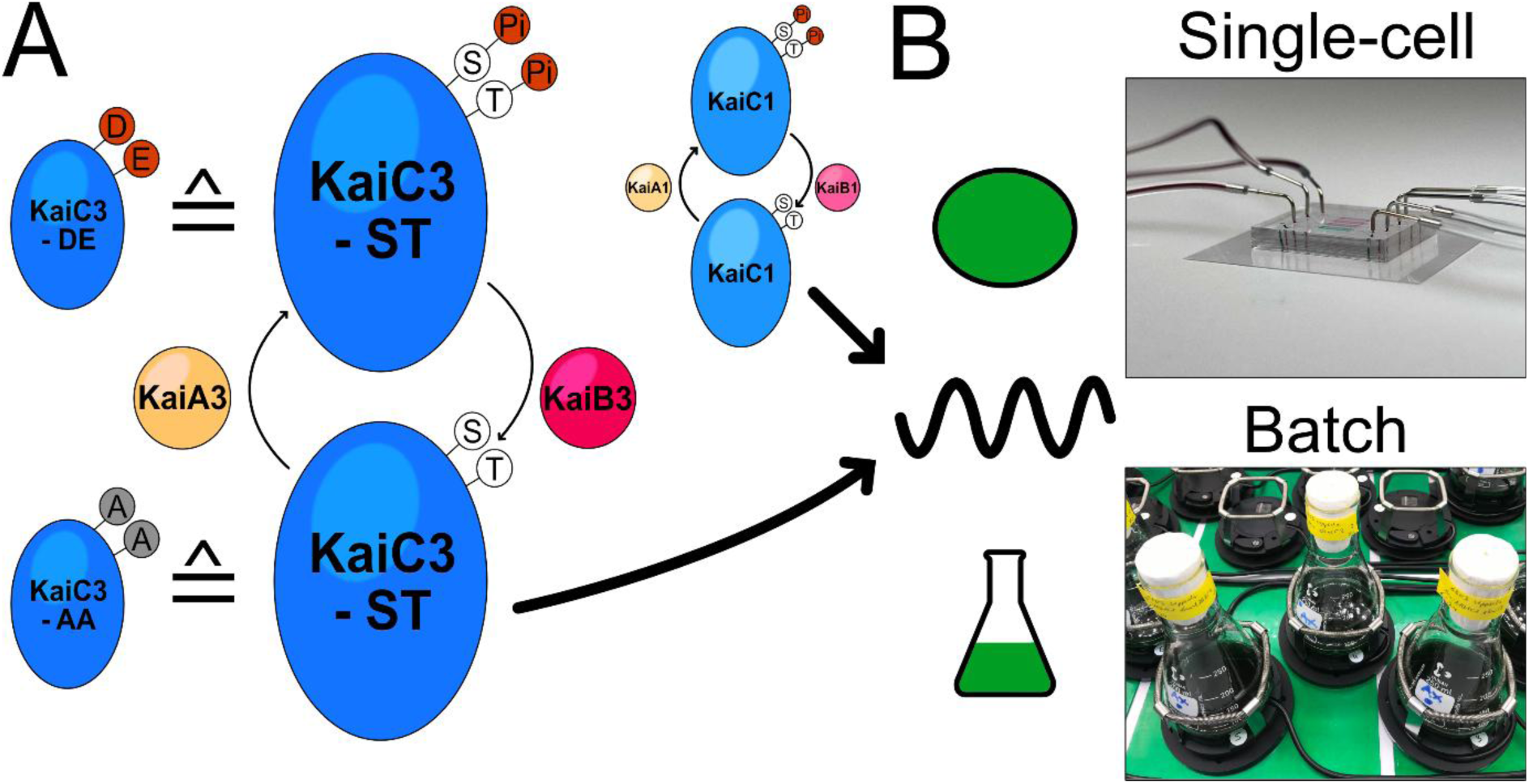
Schematic illustration of the coupled circadian oscillator in *Synechocystis*. A: Two protein complexes (KaiA1B1C1 and KaiA3B3C3) form the core of the coupled circadian oscillator. Both KaiC homologs autopphosphorylate at Serin (S) and Threonin (T) in the presence of their corresponding KaiA homolog. Binding of KaiB1 or KaiB3 initiates the dephosphorylation of KaiC1 and KaiC3, respectively. Both complexes are required for stable circadian oscillations of the backscatter [22, 34] and bioluminescence [23]. The *kaiC–DE* mutant mimics the fully phosphorylated state of KaiC3 while the *kaiC–AA* mutant mimics the fully dephosphorylated state. B: We hypothesize that both readouts (single-cell and batch) measure the same change in the refractive index of individual cells because single cells displayed circadian oscillation in phase contrast with similar parameters as backscatter oscillations.

## Discussion

### The refractive index of single cells oscillates with high robustness within a population

In this study, we used a single-cell cultivation device and time-lapse imaging to independently observe cell morphology and cell growth of *Synechocystis* under constant illumination. We demonstrated that no superimposed oscillation occurred alongside the exponential increase in colony area and that the timing of cell division was not synchronized on a 24-hour basis during single-cell cultivation. The latter observation contrasts with *Synechococcus*, where cell division is gated by the circadian clock [29, 30, 39]. Previous studies have also reported differences in the division behavior between the two strains. Martins et al. [29] observed size-like behavior in *Synechococcus*, and Yu et al. [33] found that *Synechocystis* division can best be described by an adder rule. As of today, there is no evidence for division gating in *Synechocystis,* and our investigation supports that cell division is not regulated by the clock and is not the cause of the oscillation in during batch cultivation.

We did observe an approximately 24-hour oscillation in the phase-contrast intensity of time-lapse images of *Synechocystis*. These oscillations were observed in individual cells (Fig. 1H and 2), distinct colonies as well as the mean relative phase-contrast from multiple colonies (Fig. 3D). The period and phase of the cosine approximation was similar to the period and phase measured via backscatter analysis of batch cultures, a method we established previously [34]. Therefore, we conclude that the synchronous oscillation of the refractive index of each cell leads to a cumulative backscatter signal in batch cultures. Due to the high flow rates relative to the fluid volume of the cultivation chip, putative communication between cells in different growth chambers can be neglected. Similar coherence between the circadian clocks in single cells has previously been demonstrated for *Synechococcus* [18, 25]. Knocking out *kaiC3* resulted in a loss of harmonic oscillation in the mean relative phase-contrast (Fig. 4B), thereby proving that the phase-contrast oscillation is caused by the circadian oscillator.

Fig. 1B shows that the brightness of the pixels within the single *Synechocystis* cells was heterogeneous. Rather than the overall brightness of the cells changing, it appears that the size and location of the dark spots within the cytoplasm changed periodically, leading to a 24-hour oscillation in the overall cell brightness. A plausible explanation for the oscillation is daily changes in cellular components orchestrated by the circadian oscillator. Oscillation in glycogen abundance is a likely candidate, since the deletion of circadian clock or glycogen synthesis genes abolishes the backscatter oscillation and drastically reduces glycogen levels [34]. Other potential contributors include polyhydroxybutyrate [40], DNA compaction [41–43] and polyphosphate, which has a dense crystalline structure and shows diurnal expression patterns in another cyanobacterium [44, 45]. Whether a single of these causes or a combination of them is the source of the underlying changes in the refractory properties remains to be investigated.

### The specific advantages of batch cultivation and single-cell imaging complement each other

As shown in Tab. 1, the methods presented here have specific advantages. Both methods are non-invasive. The microfluidic method enables disentangled observations of cell phenotype and cell growth under highly controlled environmental conditions, without the need for sampling or additional at-line equipment. Due to monolayer growth the light intensity can be precisely controlled and kept constant throughout the experiment. Constant perfusion of the growth medium prevents nutrient limitation. However, one image loop during single-cell cultivation lasted roughly 3–5 minutes. During this time, the lamp providing light for photosynthesis was switched off, which constrained the measurement rate to a maximum of one image per hour. Depending on the design of the microfluidic chip only a limited number of samples can be investigated in parallel. Given that an experiment takes up to 10 days, this slows down screening of different strains. On the other hand, backscatter measurements do not interrupt the cultivation process, enabling a high rate of measurement and thus better temporal resolution. While single-cell cultivation systems are often tailor-made by research groups for a defined use case the necessary equipment for backscatter analysis is commercially available. This facilitates use of the backscatter method and – depending on the purchased systems - a high number of samples can be analyzed in parallel. When used together, the two methods complement each other, enabling a more thorough investigation of the circadian clock in *Synechocystis*. Like other reporter-free methods, such as tracking daily wheel-running activity in rodents, monitoring leaf movement in plants, or observing division timing in *Synechococcus*, our method does not require genetic accessibility [1]. Similar circadian oscillation of optical properties have been observed in batch cultures of other cyanobacteria[46] Therefore, we believe our method is applicable to other cyanobacteria species and potentially to prokaryotes. We expect that our approach of combining batch and single-cell cultivation will transform circadian research by enabling scientists to investigate circadian rhythms in non-model organisms. Furthermore, our study showcases in which cases the increased effort required for single-cell cultivation is worthwhile and when researchers should opt for less labor-intensive batch cultivation.

**Table 1.** Comparison of the characteristics of the two cultivation methods presented for investigating circadian rhythms.

| Backscatter measurement | Microfluidic single-cell analysis |
| --- | --- |
| Commercially available | Tailor-made for a specific research question |
| Detection of a single parameter (backscatter) | Parallel observation of cell division, cell growth and cell morphology |
| During batch cultivation nutrients limitation can occur | Continuous perfusion of the growth medium prevents nutrient limitation |
| Measurements do not interrupt cultivation process enabling a high sampling frequency | Measurements do interrupt cultivation process limiting the sampling frequency |
| High degree of parallelization possible | Parallelization is limited by cultivation chip design and imaging frequency |

## Material and Methods

### Strains used in this study

In this study we used *Synechocystis* sp. PCC 6803 WT ‘Uppsala’ (kindly provided by Pia Lindberg, Uppsala University) and *Synechocystis* sp. PCC 6803 *ΔkaiC3* (generated in ‘Uppsala’ background [22]). Complementation strains, in which the wild-type KaiC3 (KaiC3-ST) and phosphomimetic variants KaiC3-AA and KaiC3-DE are expressed, were generated based on *Synechocystis* sp. PCC 6803 *ΔkaiC3* ‘Uppsala’ as described below.

### Generation of *ΔkaiC3* complementation strains

To re-introduce the wild-type *kaiC3* gene into *Synechocystis* sp. *ΔkaiC3,* we amplified five fragments using primers that incorporate PaqCI restriction sites (refer to Tab. S3): (1.) The plasmid backbone containing flanking regions of *kaiC3* was amplified from pJET_*dkaiC3*-Cmr [47] using primers 2431/2432. (2.) The region encoding the first 434 amino acids of KaiC3 was amplified from pASK-*kaiC3* [48] with primers 2423/2424. (3.) The remaining coding region of *kaiC3* along with 41 bp downstream of *kaiC3* were amplified from genomic DNA with primers 2425/2426. The downstream region was included, because it may function as *kaiC3* transcription termination site (see also Fig. S11). (4.) The SpecR cassette was amplified from pSHDY [49] using primers 2427/2428. (5.) The final 105 bp of *kaiC3* plus an additional 15 bp downstream were amplified from genomic DNA using primers 2429/2430, because this region may serve as promoter/RBS for the subsequent gene downstream of *kaiC3* (ssr3304, see also Fig. S11). The fragments 1-5 were assembled via Golden Gate cloning [50]. *Synechocystis ΔkaiC3* was transformed with the resultant plasmid pJET-*kaiC3*-SpecR to replace the chloramphenicol cassette with the assembled fragments, spanning from the upstream to downstream region of *kaiC3*, via homologues recombination. Complementation strains expressing the phosphomimetic variants KaiC3-DE (mimicking constant phosphorylation) and KaiC3-AA (mimicking constant dephosphorylation) were generated accordingly, using pASK*kaiC3*-DE and pASK-*kaiC3*-AA [48] as templates for step (2) described above. The genomic context is illustrated in Fig. S11. Transformants were selected using spectinomycin. Despite several rounds of segregation, some cells were able to grow on plates containing CmR, indicating that full segregation was not achieved. Nonetheless, protein levels of KaiC3 variants in the complementation strains were similar to those of native KaiC3 levels in the WT (Fig. S11).

### Microfluidic cultivation at single-cell resolution

#### Experimental setup

For microfluidic single-cell analysis, a Ti-E inverted microscope (Nikon, Japan) modified for phototrophic growth was used. For a detailed description of the cultivation setup the reader is referred to [51], and the specific cultivation setting are listed in Table 2. The intermediate magnification lens was set to 1.5x to further enhance the magnification in images of *Synechocystis*. The microscope was controlled using NIS-Elements 4.51.

**Table 2:** Device setting during microfluidic single-cell cultivation.

| Device | Product name (Supplier) | Specific setting |
| --- | --- | --- |
| Objective | Plan Apo $\lambda$ 100x Oil Ph3 DM, NA = 1.45 (Nikon, Japan) | - |
| Temperature incubator and controller | Incubator NL 2000, Temp-Controller 2000-2 (PECON, Germany) | Temperature at 30 °C |
| Growth light | Spectra Tune Lab (LED MOTIVE, Spain) | 4 $\mu\text{mol}\cdot\text{m}^{-2}\cdot\text{s}^{-1}$ of artificial sunlight (Fig. S1) |
| Syringe pump | Nemesys (CETONI, Germany) | Volumetric flow rate (BG11) = 200 $\text{nL}\cdot\text{min}^{-1}$ |
| Camera | DS-Fi3 (Nikon, Japan) | Color disabled |

### Precultivation

Precultivation of cyanobacteria was conducted in a MC-1000 OD Multi-Cultivator (Photon Systems Instruments; Czech Republic) for approximately 24 hours. The light-intensity was set to 80 µmol_(Photons)_·m^-2^·s^-1^ cool white light. The cyanobacteria were inoculated to an OD_720_ of 0.1 in 50 mL BG11 growth medium. The device was programmed to perform a 12h/12h day/night cycle. Samples were taken shortly after the end of the night.

### Time-lapse microscopy

The master mold for PDMS soft lithography molding is made up of a 4’’ silicon wafer with two layers of SU-8 photoresist. It was produced using mask-based photolithography at the Helmholtz Nano Facility [52]. For soft lithography, PDMS and curing agent (SYLGARD™184 Silicone Elastomer Kit) were mixed at a ratio of 10:1, degassed and poured over the master mold. The mixture was exposed to 80 °C for 2h, afterwards the polymerized PDMS slab was removed and cut into appropriate pieces. Chips were prepared in bulk and stored until use.

For the final assembly the cultivation chip (Fig. 7) and a glass substrate (D263®Bio, 39.5 mm × 34.5 mm × 0.175 mm; Schott AG, Germany) were exposed to oxygen plasma for 25 s. The side of the cultivation chip containing the cavities was flipped onto the glass substrate to initiate covalent bonding. This step was conducted the day before the experiment began. For a more detailed description of the molding and assembly of the PDMS cultivation chips, the reader is referred to previous publications [51, 53].

**Figure 7:**
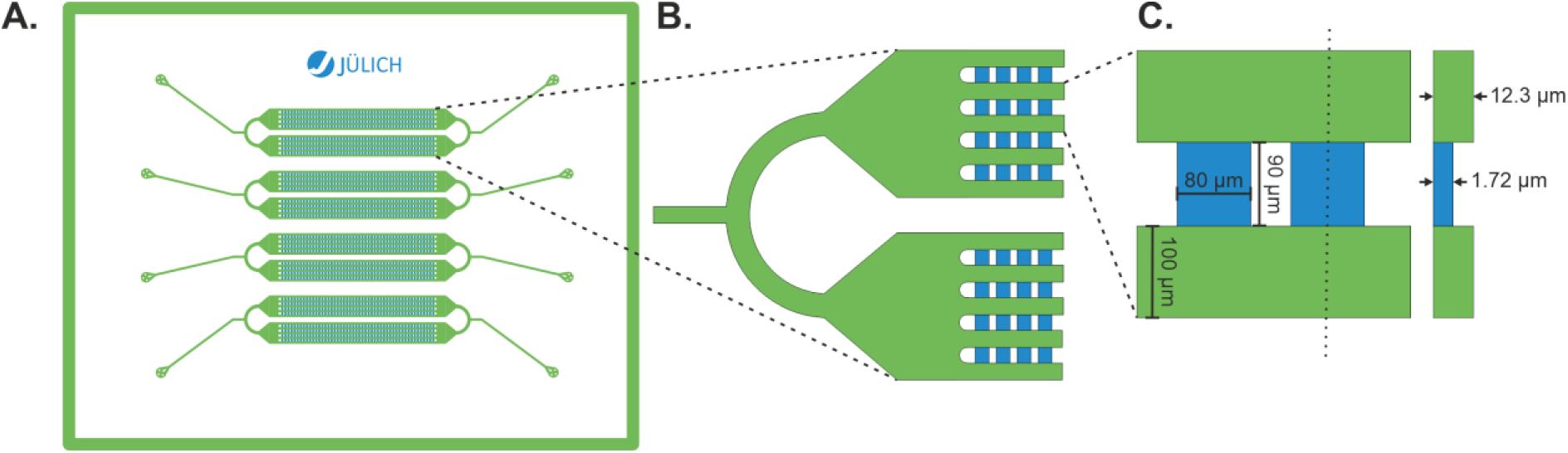
Cultivation layout used for microfluidic single-cell analysis. A: The cultivation layout includes four separate cultivation arrays. B: Each array includes 8 rows of 50 chambers, that are interconnected by shared supply channels with a height of 10 µm. C: The growth chambers are 80 x 90 μm in size and 1.72 μm high.

The cultivation chips were fixed on the microscope stage using a conventional transparent adhesive tape. The incubator was heated overnight to prevent the loss of focus owing to the thermal expansion of the PDMS. Prior to starting the experiment, the syringes (Omnifix-F 3 mL, Braun, Germany) containing the growth medium were connected to the cultivation chips. The cultivation chips were flushed with the medium for at least 30 min. The cells were then inoculated into the cultivation chip using a syringe (Omnifix-F 1 mL, Braun, Germany) and a small piece of tubing. For each condition, 10-15 growth chambers, each containing one centrally placed cell, were manually selected, before the experiment was started. During time-lapse microscopy phase-contrast and fluorescence images (Ex: 514/30 nm; Dm: 561 nm; Em: 629/56 nm) were captured every hour.

The two cultivation methods were performed in different laboratories using slightly different BG11 recipes. The backscatter lab used 10 mM TES buffer at a pH of 8, whereas the microfluidic lab used 10 mM HEPES buffer at a pH of 7.5. Additionally, the final concentration of EDTA-Na₂ was 1 mg/L in the microfluidic lab and 0.521 mg/L in the backscatter laboratory. As a control the WT was analyzed again in both BG11 variants using the microfluidic device and no difference in the growth behavior was observed. Underlying data (Fig. S8, Tab. S2) and a comparison of the medium recipes (Tab. S1) is shown in the Supplementary Material.

### Post-experimental procedure

The post-experimental procedure has been previously explained in detail [51]. The process included the following steps: (i) Drift correction, rotation, and cropping in Fiji [54]. (ii) Uploading of the image sequences to the OMERO [55] server. (iii) Subsequent deep-learning cell segmentation using a Python script [35]. During the segmentation process, data on the single-cell area, cultivation time, autofluorescence, and phase-contrast intensity were gathered for each segmented cell. Multiple analyses were performed on these data tables.

#### Division timing

For each image sequence, an array containing information on the number of cells over time was formed. The cell number data only contain positive integers. A Python script looped through each array and checked if the current value was greater than the value in the previous row. If so, a division event occurred, and the corresponding time was added to the list of all division events. After looping through all the arrays, the subjective time was calculated from the cultivation time, and the data were plotted as a histogram.

#### Growth rate

For each image sequence, an array was formed that contained information on the cumulative cell area (colony area) over the cultivation time. Data from the first 24 h was omitted. The growth rate was calculated according to an exponential model by performing a linear fit on the natural logarithm of the colony area. The slope of the linear fit represents the growth rate of the colony.

#### Phase contrast oscillation

For each image sequence, an array was formed containing information on the average phase-contrast intensity over the cultivation time. To normalize the data, a Python-based workflow adapted from [34] was used. The workflow included the following steps

- Fitting a sixth-degree polynomial through the phase-contrast intensity data.
- Calculation of the residuals by subtracting the raw data from the polynomial fit
- Smoothing the data by applying a six-point moving average to the signal.
- Calculating the mean and standard deviation from multiple image sequences off the same growth array.

To evaluate and compare parameters of the oscillation between different experiments a DFT was performed on the mean phase-contrast signal excluding the first 24 h using NumPy [56]. Due to the broad width of the frequency bins resulting from the limited number of samples in microfluidic analyses, the dominant cosine wave was further optimized using the dominant frequency, corresponding phase, and amplitude as initial parameters. Then, the resulting parameters for amplitude, phase and period were compared between strains and experiments.

### Batch cultivation with backscatter measurement

#### Precultivation

The cultures were first inoculated by suspending cell material from BG11-agar plates in 20 ml of BG11 in 100 ml shake flasks. These first precultures were incubated for 7 days (30 °C, 150 rpm shaking, 0.5 % CO_2_, 50 % humidity, 32 µmol_(photons)_·m^-2^·s^-1^) in a 257LX1 incubator (Labwit, Australia) after which they reached an OD_750nm_ between 2.4 and 4.6. These cultures were diluted by adding 80 ml BG11 to a final volume of 100 ml in 250 ml shake flasks and incubated for an additional 3 days. Before the final dilution and the start of the experiment, the cultures reached an OD_750nm_ of 1.7 to 2.2.

### Backscatter measurement

To start the experiment, the second precultures were split and diluted to an OD_750nm_ of 1.0 with 2x BG11 to a final volume of 50 ml. These cultures were incubated in an Multitron HT Incubator (Infors, Switzerland) (30 °C, 150 rpm shaking, 1 % CO2, 50 % humidity, 32 µmol_(photons)_·m^-2^·s^-1^ (The spectrum is shown in Fig. S1)) for up to ten days. The Incubators were equipped with 16 Cell Growth Quantifier sensors (Scientific Bioprocessing, Germany) which measured the backscatter of the cultures every 20 seconds at a wavelength of 730 nm. For a more detailed description, see Berwanger et al. [34].

### Data analysis

To isolate the circadian oscillation from the raw backscatter measurements, a 5th-degree polynomial was fitted to the data. The polynomial represented only the growth component of the raw data, excluding any oscillation signal. To extract the signal, the polynomial was subtracted from the raw data. The residuals were then normalized by subtraction of the arithmetic mean and smoothed by a rolling average. For more details, see [34]. For graphical representation, the arithmetic means of all replicates of one strain in one experiment are shown. The smoothed signal was used for further analysis. To determine the period, amplitude and phase of the oscillation, a discrete Fourier transformation using the fast Fourier algorithm was performed on the smoothed signal. From the resulting frequency spectra, we calculated the period by taking the reciprocal of the most dominant frequency peak.

### Uncertainty calculation for backscatter

Because the number of replicates for each backscatter experiment was limited, only two to three replicates for each strain were measured per experiment. Therefore, we displayed the uncertainty of the measurement as the 95 % confidence interval of a Student’s t-distribution. The arithmetic mean, standard deviation, standard error of the mean and 95 % confidence intervals were calculated using the following formulas respectively:

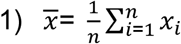

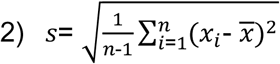

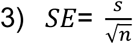

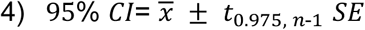

Where *x̄* is the arithmetic mean, *s* the standard deviation, *SE* the standard error of the mean, 95 % CI the 95 % confidence interval and *n* the number of replicates.

### Discrete Fourier transformation

The Discrete Fourier Transformation (DFT) can be used to convert a discrete, time-dependent signal x_n_ comprising N samples into the frequency domain. The result is a complex array X_k_.

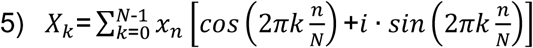

The fast Fourier transformation algorithm is commonly applied to efficiently calculate the DFT from a time-dependent signal with a large number of samples. The frequencies corresponding to the values of k can be calculated with the sampling interval *Δt* using equation 6.

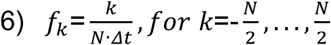

Note that f_k_ includes negative frequencies. However, for real inputs of x_n_, the spectrum is symmetrical, hence *X[f] = X[-f]*. The frequency at which the value of X_f_ is at its maximum is referred to as the dominant frequency. In order to reconstruct the cosine wave at the dominant frequency (f_dom_ [h^-1^]) the corresponding period (P [h]), the dominant amplitude (A [-]) and phase (φ [h]) can be calculated as follows.

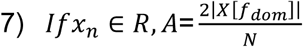

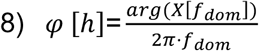

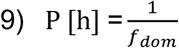

Equ. 10 was used to convert the phase (φ [h]) into the time at which the first peak of the fitted cosine wave occurred.

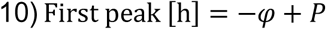

The dominant cosine wave is then expressed as [57]:

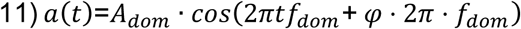

## Acknowledgments

We would like to thank Sabrina Zander from the DataPLANT team for her help setting up the ARC data repository as well as Johannes Seiffarth help with troubleshooting the acia-based automated cell segmentation and data analysis workflows. We thank Markus Kollmann for his support.

## Generative AI statement

The authors used ChatGPT Model 5 and Claude Sonnet 4.5 to assist with generating Python code for analyzing circadian rhythms. All AI-generated code was manually inspected and corrected as necessary before implementation. Additionally, the authors used DeepL and PaperPal to refine pre-written manuscript text. All text passages were manually checked for accuracy and edited as needed

## Data Availability Statement

All data generated for this study is deposited in the PLANTdataHUB [58]: (URL: https://doi.org/10.60534/9j7k0-8b423)

## Author contribution

**Conceptualization:** Ilka M. Axmann and Anika Wiegard

**Formal analysis:** Lennart Witting and Florian P. Stirba

**Funding acquisition:** Ilka M. Axmann and Dietrich Kohlheyer

**Investigation:** Lennart Witting, Florian P. Stirba, Anika Wiegard, Ekaterina Ivanova, and Petra Kolkhof

**Methodology:** Lennart Witting, Florian P. Stirba and Julius Nohr

**Software:** Julius Nohr, Florian P. Stirba, und Lennart Witting

**Supervision:** Ilka M. Axmann, Anika Wiegard, and Dietrich Kohlheyer

**Visualization:** Lennart Witting and Florian P. Stirba

**Writing – original draft:** Lennart Witting, Florian P. Stirba, and Anika Wiegard

**Writing – review & editing:** Anika Wiegard and Ilka M. Axmann

## Supporting information

**Figure S1: Spectral distribution of the light used during single-cell and batch cultivation.** A: For single-cell cultivation, the Spectra Tune Lab light engine (LEDMOTIVE, Spain) was set to emit blackbody radiation at 5,800 K to mimic the spectrum emitted by the sun. Light intensity was measured directly under the cultivation chip using an LI190R quantum sensor (LI-COR, USA) to ensure precise illumination. B: The spectrum of the built-in lamp of the incubator was measured using an AvaSpec-Mini spectrometer and AvaSoft software version 8.16.1.0 (Avantes, Netherlands). Measurements were conducted with the sensor placed in the middle of the incubator, and the dark incubator was used as a blank. The integration time was 2,600 ms.

**Figure S2: Replicate experiment to Fig. 4.** The replicate experiment was performed after the experiment shown in Fig. 4. with bacteria from a different plate but stemming from the same frozen culture stock. A-C: Relative phase-contrast intensity and dominant cosine wave for the WT (blue; same data as in Fig. S7 A), Δ*kaiC3* knockout (orange), and *kaiC3*-ST complementation mutant (purple). Note that the initial sharp increase in phase-contrast intensity observed in A-C was caused by a loss of focus during the first hours of the time-lapse experiment. These were always excluded from the DFT analysis. D - E: Boxplots displaying the growth rates derived from colony area plots (D) and single-cell areas (E) during single-cell cultivation for the WT, Δ*kaiC3* knockout, and *kaiC3*-ST complementation mutant. The growth rates were compared by one-way ANOVA followed by Tukey’s HSD test (* p < 0.05).

**Figure S3: Raw single-cell phase-contrast intensity data for the Δ*kaiC3* knockout mutant.** Each plot contains data from the time-lapse image sequence of a single colony acquired during a continuous microfluidic experiment. Although each colony was located in a separate growth chamber, all growth chambers belonged to the same array of chambers on the microfluidic chip. Therefore, they are interconnected through shared supply channels. Each data point represents the mean intensity of all pixels within the boundaries of an individual *Synechocystis* cell.

**Figure S4: Raw single-cell phase-contrast intensity for the *kaiC3*-ST complementation mutant.** Each plot contains data from the time-lapse image sequence of a single colony acquired during a continuous microfluidic experiment. Although each colony was located in a separate growth chamber, all growth chambers belonged to the same array of chambers on the microfluidic chip. Therefore, they are interconnected by shared supply channels. Each data point represents the mean intensity of all pixels within the boundary of an individual *Synechocystis* cell.

**Figure S5: Oscillation parameters calculated from each image sequence individually.** In Figs 3 and 4, the DFT was performed on the mean relative phase-contrast of multiple colonies to demonstrate synchrony. For this figure, DFT and subsequent cosine fitting were performed separately for each colony/image sequence. The fact that the single-colony periods for Δ*kaiC3* do not cluster around 24 h, as they do for WT and *kaiC3*-ST, shows that knocking out the *kaiC3* gene dampens the oscillation itself rather than just abolishing the synchrony between colonies.

**Figure S6: Colony area over time for WT, Δ*kaiC3* knockout mutant, and *kaiC3*-ST complementation mutant.** For each strain, the mean and standard deviation of n colonies are shown.

***Figure S7:* Growth of the WT at 7 µmol(photons)·m^-2^·s^-1^ compared to growth at 4 µmol(photons)·m^-2^·s^-1^.** A: Mean relative phase-contrast intensity of n independent colonies and the dominant cosine wave for growth at 7 µmol(photons)·m^-2^·s^-1^ (blue; same data as Fig. 4 WT) and 4 µmol(photons)·m^-2^·s^-1^ (green). The differences in period (blue: 24.63 h; green: 24.88 h), phase (blue: −0.17 h; green: 2.71 h), and amplitude (blue: 2.24 a.u.; green: 0.75 a.u.) of the oscillation between light intensities did not exceed the differences observed between independent experiments at the same light intensity. B: Mean colony area of n independent colonies. At 7 µmol(photons)·m^-2^·s^-1^ growth of the colonies was faster than at 4 µmol(photons)·m^-2^·s^-1^ (Same data as Fig. S6). C: Growth rates derived from colony area measurements. The growth rates (C) were compared by one-way ANOVA followed by Tukey’s HSD test (⁕ p < 0.05). D: Single-cell areas observed at different light intensities.

**Table S1: BG11 recipe comparison.** Differences are highlighted in bold.

**Figure S8: Growth of the *Synechocystis* WT during single-cell cultivation in the microfluidic photobioreactor for both BG11 recipes.** For the BG11 recipe used in the single-cell cultivation laboratory (A), the mean colony area and mean relative phase contrast were calculated from 15 independent colonies. For the recipe from the batch cultivation laboratory (B) the mean of 14 colonies is shown. The growth rates (C) were compared by one-way ANOVA followed by Tukey’s HSD test (⁕ p < 0.05). The single cell areas are shown in D.

**Table S2: Comparison of cosine fit parameters for different BG11 recipes.** The results stem from a single single-cell cultivation experiment in which two growth channels were inoculated with WT. One of these arrays was perfused with BG11 from the single-cell cultivation laboratory. The other was perfused with BG11 from the batch cultivation laboratory.

**Figure S9: Raw backscatter measurements for the individual replicates** of the strains WT (blue), Δ*kaiC3* (orange), *kaiC3*-ST (purple), *kaiC3*-AA (green) and *kaiC3*-DE (red).

**Figure S10: Isolated smoothed backscatter oscillation for the individual replicates** of the strains WT (blue), Δ*kaiC3* (orange), *kaiC3*-ST (purple), *kaiC3*-AA (green) and *kaiC3*-DE (red).

**Figure S11: Genomic context of *kaiC3* and generation of complementation strains.** A: According to the annotation of Mitschke et al., a TATA box (turquoise) is present at the end of the coding sequence (CDS) of *kaiC3* (blue) and the 5’UTR (violet) of the hypothetical protein Ssr3304 starts directly downstream of *kaiC3*, followed by the CDS of Ssr3304 (grey, 38 bp downstream of *kaiC3*) [1]. Accordingly, the region downstream of *kaiC3* may function as a terminator for *kaiC3* transcription, and the end of the *kaiC3* CDS may also function as a promoter/RBS for ssr3304. B: In *Synechocystis* sp. PCC 6803 Δ*kaiC3* a CmR resistance cassette, together with a region of a plasmid, was inserted between the upstream and downstream flanking regions of k*aiC3* [2,3]. C: In the *kaiC3* complementation strains, the CmR cassette in *Synechocystis* sp. PCC6803 Δ*kaiC3* [3] was exchanged with the respective *kaiC3* coding sequences (*kaiC3*-ST, *kaiC3*-AA, or *kaiC3*-DE). To maintain the native terminator of *kaiC3*, we kept the 38bp downstream region of kaiC3 and added a SpecR cassette as a selection marker behind it. We inserted the last 105 bp of *kaiC3* behind the SpecR cassette to maintain the hypothetical promoter of ssr3304 intact. Genomic maps were generated using the SnapGene software. D*: Synechocystis* sp. PCC 6803 WT and complementation strains (Δ*kaiC3* +*kaiC3*-AA, Δ*kaiC3* +*kaiC3*-DE, Δ*kaiC3*+*kaiC3*) were inoculated in 20 ml BG11 and grown at 150 rpm, 0.5 % CO2, and constant illumination with approximately 25 µmol(photons)·m^-2^·s^-1^. On days 5 and 6, cells were diluted to an OD750nm of 0.4 and grown for two more days to OD750nm ∼1.3-1.5. Cells from 4 ml culture were sheared with glass beads in lysis buffer (8M urea, 20mM HEPES pH 8.0). Whole cell extracts were normalized to 100 ng Chl a content, separated via SDS-PAGE using a 7.5% PAA gel, and subsequently blotted and immunodecorated with a KaiC3-specific antibody (1:3.750 in TBST) [4] and goat anti-rabbit IgG (H + L) Secondary Antibody, HRP (Thermo Fisher Scientific, 1:50,000, 1 h RT). After applying the Pierce SuperSignal West Pico detection reagent (Thermo Scientific), the signal was imaged using a ChemiDoc XRS+ Imaging System with ImageLab^TM^ Software (BioRad).

**Table S3: List of primers used to generate the investigated strains.** The PaqCI recognition site (cacctgc) is indicated in bold. PaqCI cleaves the leading strand four bp downstream of the recognition site and the lagging strand eight bp downstream of the recognition site. In the table, the four bases following the recognition sequence are marked in italics. The complementary sticky ends produced after cleavage are marked in color, bold, and underlined. Primers were designed using the NEBridge Golden Gate assembly tool from New England biolabs.

