## Supplement for "Live-cell imaging enables reporter-free monitoring of the circadian rhythm in individual *Synechocystis* cells"

### Spectral power distribution

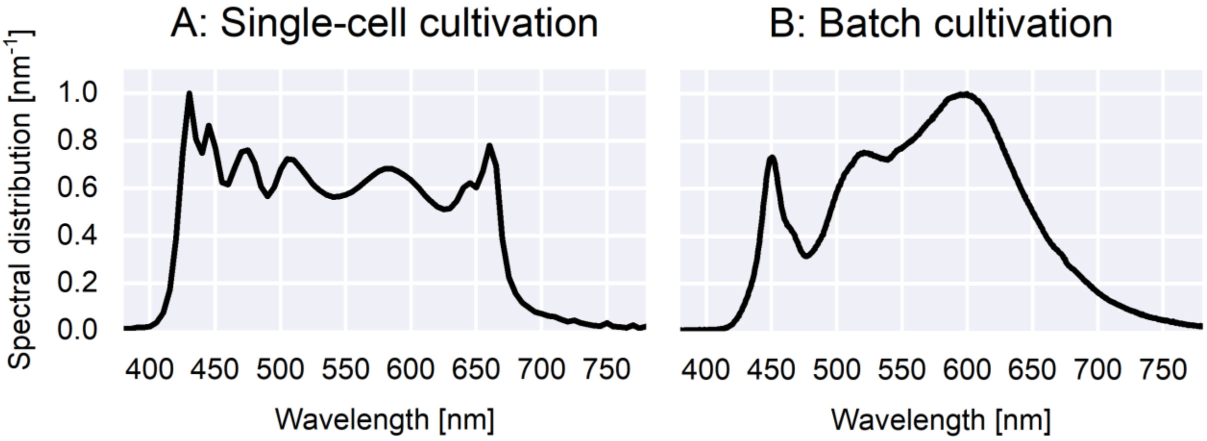


**Figure S1: Spectral distribution of the light used during single-cell and batch cultivation.** A: For single-cell cultivation, the Spectra Tune Lab light engine (LEDMOTIVE, Spain) was set to emit blackbody radiation at 5,800 K to mimic the spectrum emitted by the sun. Light intensity was measured directly under the cultivation chip using an LI190R quantum sensor (LI-COR, USA) to ensure precise illumination. B: The spectrum of the built-in lamp of the incubator was measured using an AvaSpec-Mini spectrometer and AvaSoft software version 8.16.1.0 (Avantes, Netherlands). Measurements were conducted with the sensor placed in the middle of the incubator, and the dark incubator was used as a blank. The integration time was 2,600 ms.

### Supplementary microfluidic data

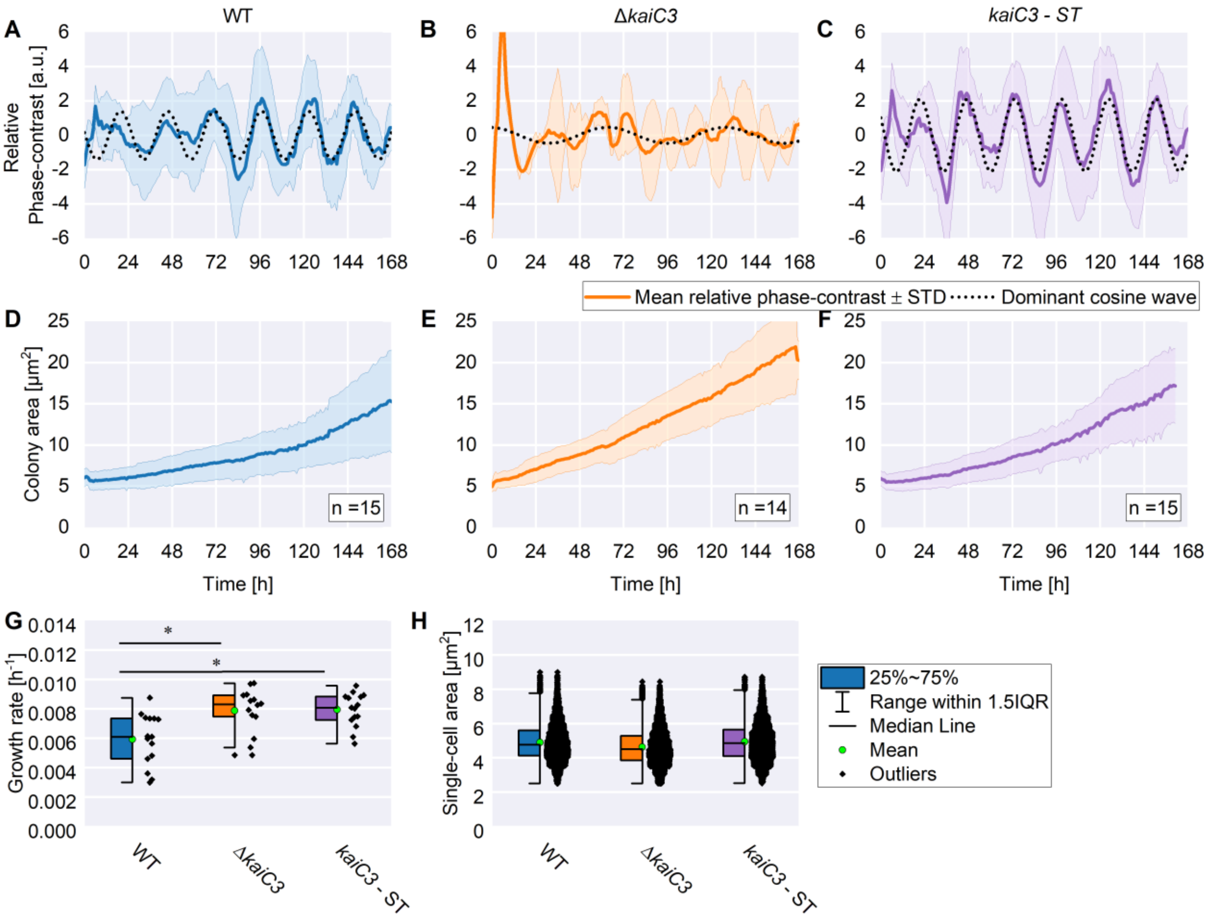


**Figure S2: Replicate experiment to Fig. 4.** The replicate experiment was performed after the experiment shown in Fig. 4. with bacteria from a different plate but stemming from the same frozen culture stock. A-C: Relative phase-contrast intensity and dominant cosine wave for the WT (blue; same data as in Fig. S7 A), *ΔkaiC3* knockout (orange), and *kaiC3*-ST complementation mutant (purple). Note that the initial sharp increase in phase-contrast intensity observed in A-C was caused by a loss of focus during the first hours of the time-lapse experiment. These were always excluded from the DFT analysis. D - E: Boxplots displaying the growth rates derived from colony area plots (D) and single-cell areas (E) during single-cell cultivation for the WT, Δ*kaiC3* knockout, and *kaiC3*-ST complementation mutant. The growth rates were compared by one-way ANOVA followed by Tukey's HSD test (* $p < 0.05$).

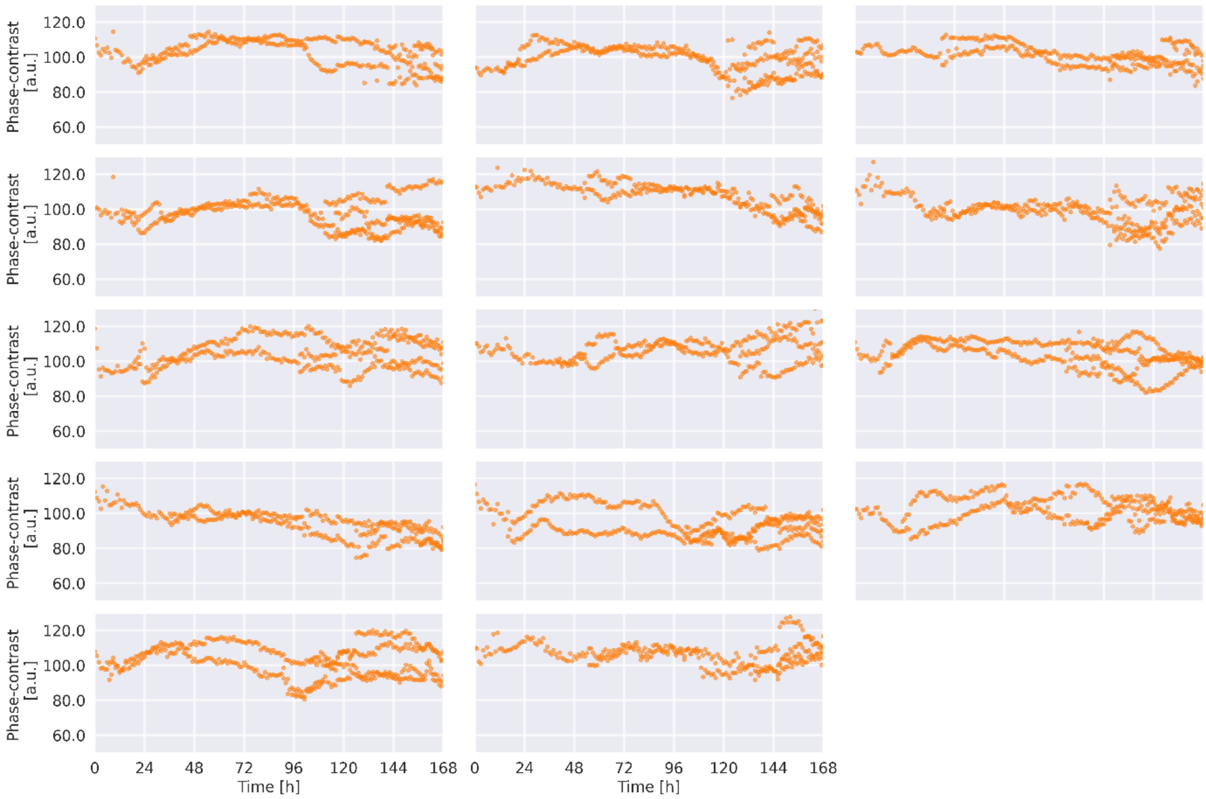


**Figure S3: Raw single-cell phase-contrast intensity data for the Δ*kaiC3* knockout mutant**. Each plot contains data from the time-lapse image sequence of a single colony acquired during a continuous microfluidic experiment. Although each colony was located in a separate growth chamber, all growth chambers belonged to the same array of chambers on the microfluidic chip. Therefore, they are interconnected through shared supply channels. Each data point represents the mean intensity of all pixels within the boundaries of an individual *Synechocystis* cell.

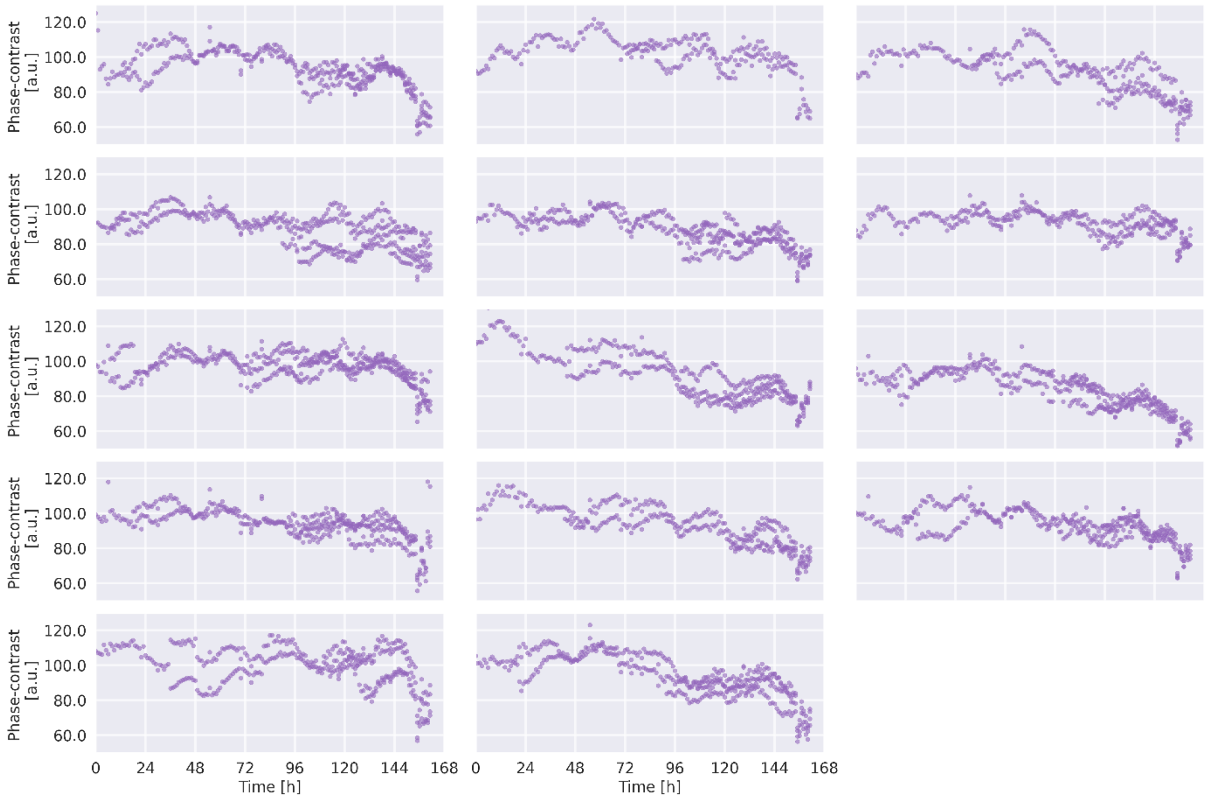


**Figure S4: Raw single-cell phase-contrast intensity for the *kaiC3*-ST complementation mutant.** Each plot contains data from the time-lapse image sequence of a single colony acquired during a continuous microfluidic experiment. Although each colony was located in a separate growth chamber, all growth chambers belonged to the same array of chambers on the microfluidic chip. Therefore, they are interconnected by shared supply channels. Each data point represents the mean intensity of all pixels within the boundary of an individual *Synechocystis* cell.

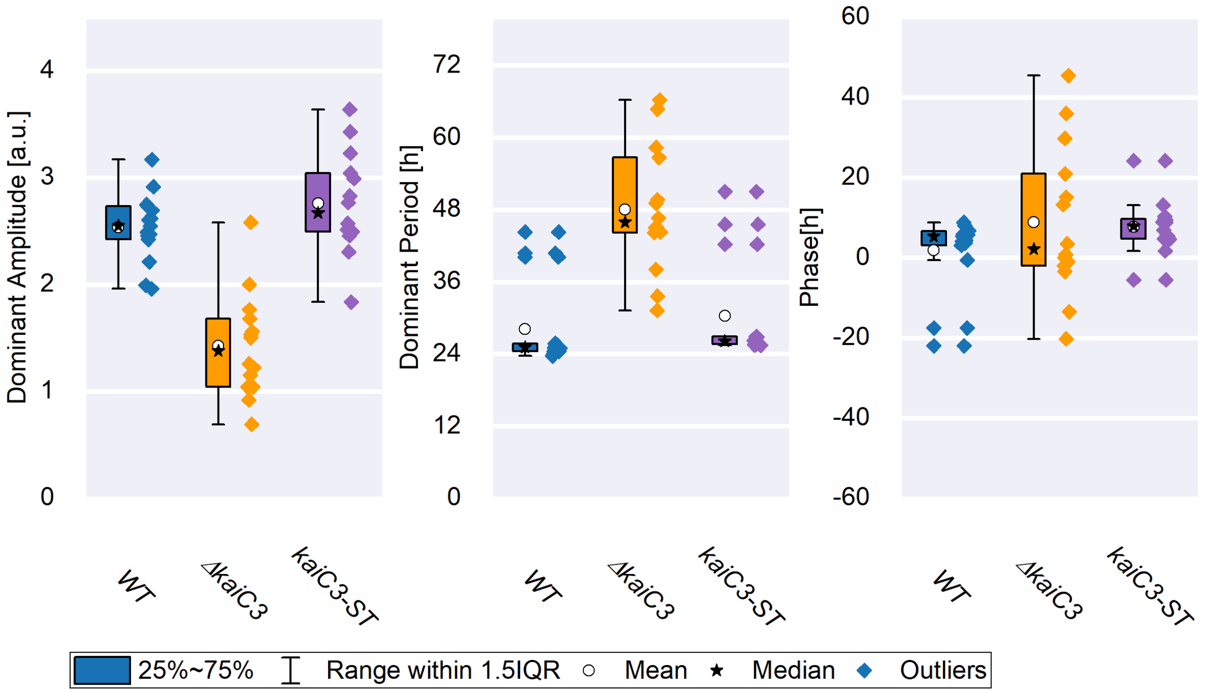


**Figure S5: Oscillation parameters calculated from each image sequence individually.** In Figs 3 and 4, the DFT was performed on the mean relative phase-contrast of multiple colonies to demonstrate synchrony. For this figure, DFT and subsequent cosine fitting were performed separately for each colony/image sequence. The fact that the single-colony periods for *ΔkaiC3* do not cluster around 24 h, as they do for WT and *kaiC3*-ST, shows that knocking out the *kaiC3* gene dampens the oscillation itself rather than just abolishing the synchrony between colonies.

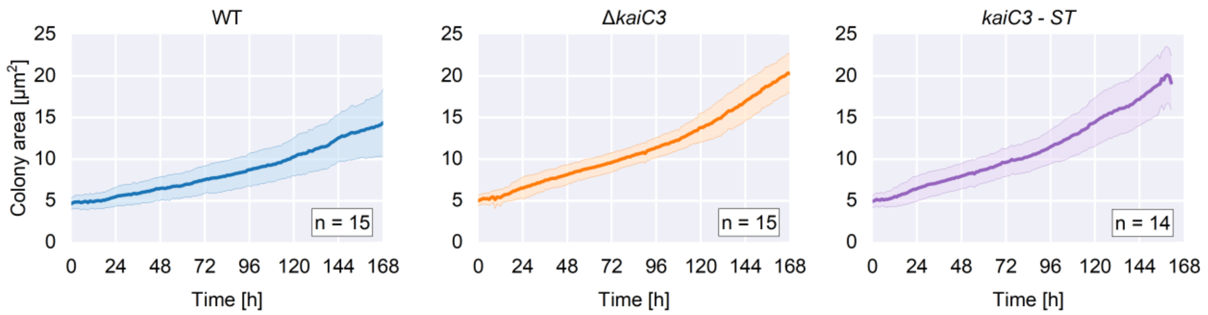


**Figure S6: Colony area over time for WT, *ΔkaiC3* knockout mutant, and *kaiC3*-ST complementation mutant.** For each strain, the mean and standard deviation of n colonies are shown.

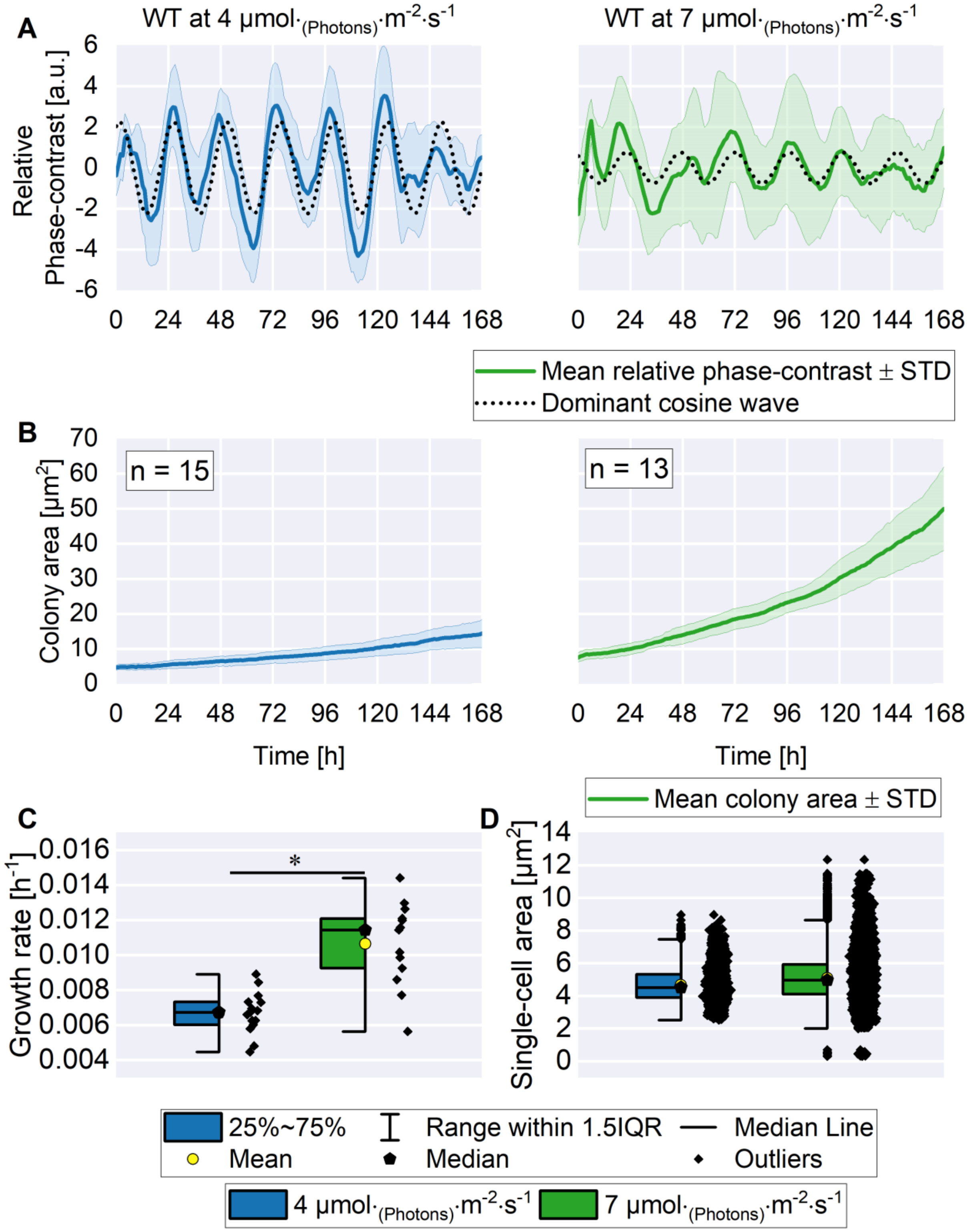


**Figure S7: Growth of the WT at 7 µmol$_{(photons)}$·m$^{-2}$·s$^{-1}$ compared to growth at 4 µmol$_{(photons)}$·m$^{-2}$·s$^{-1}$.** A: Mean relative phase-contrast intensity of n independent colonies and the dominant cosine wave for growth at 7 µmol$_{(photons)}$·m$^{-2}$·s$^{-1}$ (blue; same data as Fig. 4 WT) and 4 µmol$_{(photons)}$·m$^{-2}$·s$^{-1}$ (green). The differences in period (blue: 24.63 h; green: 24.88 h), phase (blue: -0.17 h; green: 2.71 h), and amplitude (blue: 2.24 a.u.; green: 0.75 a.u.) of the oscillation between light intensities did not exceed the differences observed between independent experiments at the same light intensity. B: Mean colony area of n independent colonies. At 7 µmol$_{(photons)}$·m$^{-2}$·s$^{-1}$ growth of the colonies was faster than at 4 µmol$_{(photons)}$·m$^{-2}$·s$^{-1}$ (Same data as Fig. S6). C: Growth rates derived from colony area measurements. The growth rates (C) were compared by one-way ANOVA followed by Tukey's HSD test (∗ $p < 0.05$). D: Single-cell areas observed at different light intensities.

### Comparison of BG11 medium recipes

**Table S1: Comparison of BG11 recipes.** Differences are highlighted in bold.

| Component | Concentration Single-cell cultivation laboratory | Concentration Batch cultivation laboratory |
|---|---|---|
| **$NaNO_3$** | 1.496 g/L | 1.496 g/L |
| **$MgSO_4$ x 7 $H_2O$** | 75 mg/L | 74.9 mg/L |
| **CaCl2 x 2 $H_20$** | 36 mg/L | 36 mg/L |
| **Citric acid** | 6.5 mg/L | 6 mg/L |
| **EDTA-$Na_2$** | **1 mg/L** | **0.521 mg/L** |
| **$H_3BO_3$** | 2.86 mg/L | 2.86 mg/L |
| **$MnCl_2$ x 4 $H_20$** | 1.81 mg/L | 1.81 mg/L |
| **$ZnSO_4$ x 7 $H_20$** | 0.222 mg/L | 0.222 mg/L |
| **$Na_2MoO_4$ x 2 $H_20$** | 0.391 mg/L | 0.390 mg/L |
| **$CuSO_4$ x 5$H_20$** | 0.079 mg/L | 0.079 mg/L |
| **Co($NO_3$)2 x 6 $H_20$** | 0.04947 mg/L | 0.049 mg/L |
| **$K_2HPO_4$** | 30.48 mg/L | 30 mg/L |
| **$NaCO_3$** | 20.03 mg/L | 20 mg/L |
| **Fe-$NH_4$-Citrat** | 6 mg/L | 6 mg/L |
| **Buffer** | **20 mM HEPES-NaOH pH 7.5** | **10 mM TES pH 8** |

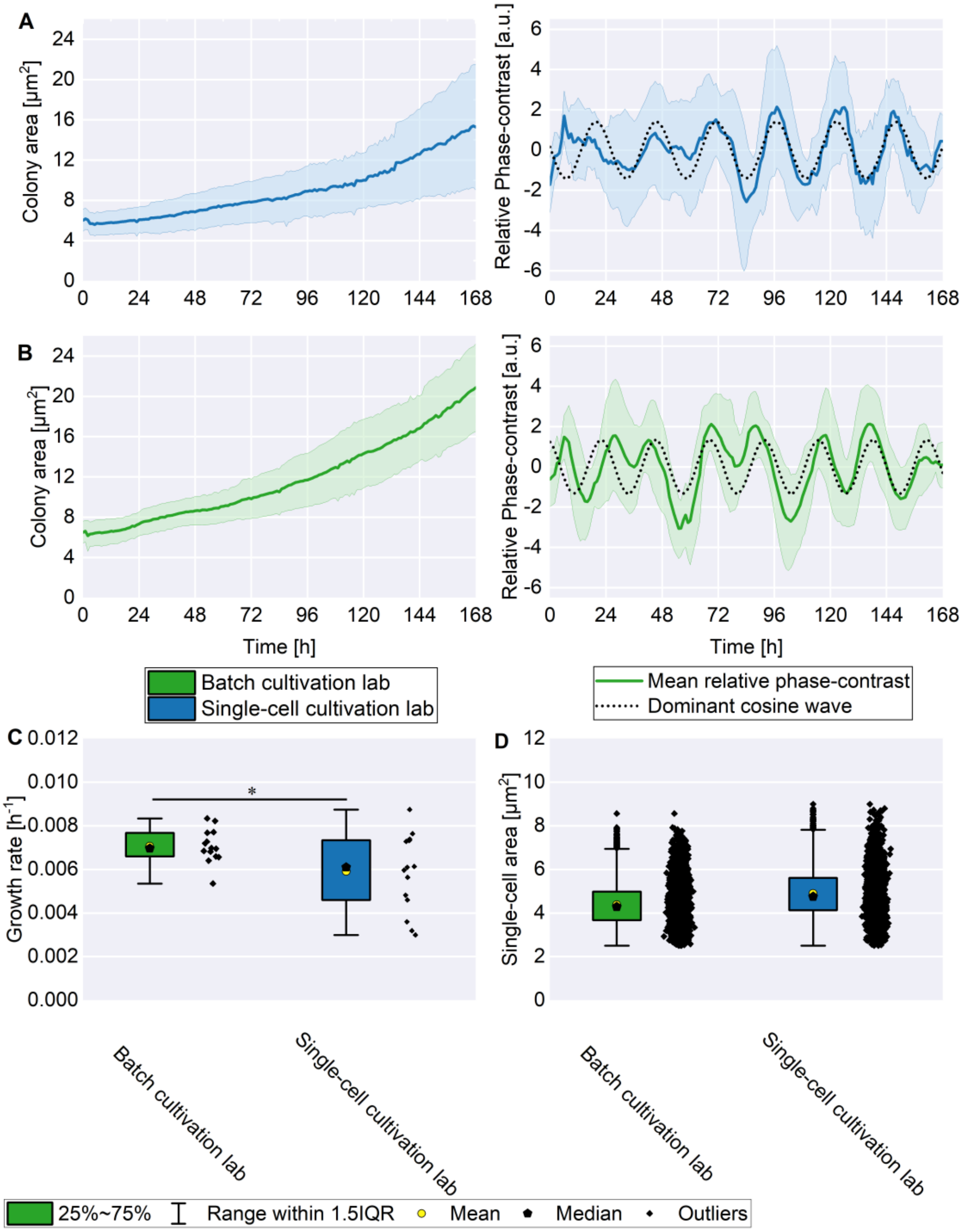


**Figure S8: Growth of the *Synechocystis* WT during single-cell cultivation in the microfluidic photobioreactor for both BG11 recipes**. For the BG11 recipe used in the single-cell cultivation laboratory (A), the mean colony area and mean relative phase contrast were calculated from 15 independent colonies. For the recipe from the batch cultivation laboratory (B) the mean of 14 colonies is shown. The growth rates (C) were compared by one-way ANOVA followed by Tukey's HSD test (∗ $p < 0.05$). The single cell areas are shown in D.

**Table S2: Comparison of cosine fit parameters for different BG11 recipes.** The results stem from a single single-cell cultivation experiment in which two growth channels were inoculated with WT. One of these arrays was perfused with BG11 from the single-cell cultivation laboratory. The other was perfused with BG11 from the batch cultivation laboratory.

| BG11 recipe | Period [h] | Amplitude [-] | Phase [h] | First Peak [h] |
|---|---|---|---|---|
| Batch cultivation laboratory | 23.24 | 1.33 | 1.37 | 21.87 |
| Single-cell cultivation laboratory | 25.69 | 1.41 | 5.97 | 19.72 |

### Supplementary batch cultivation data

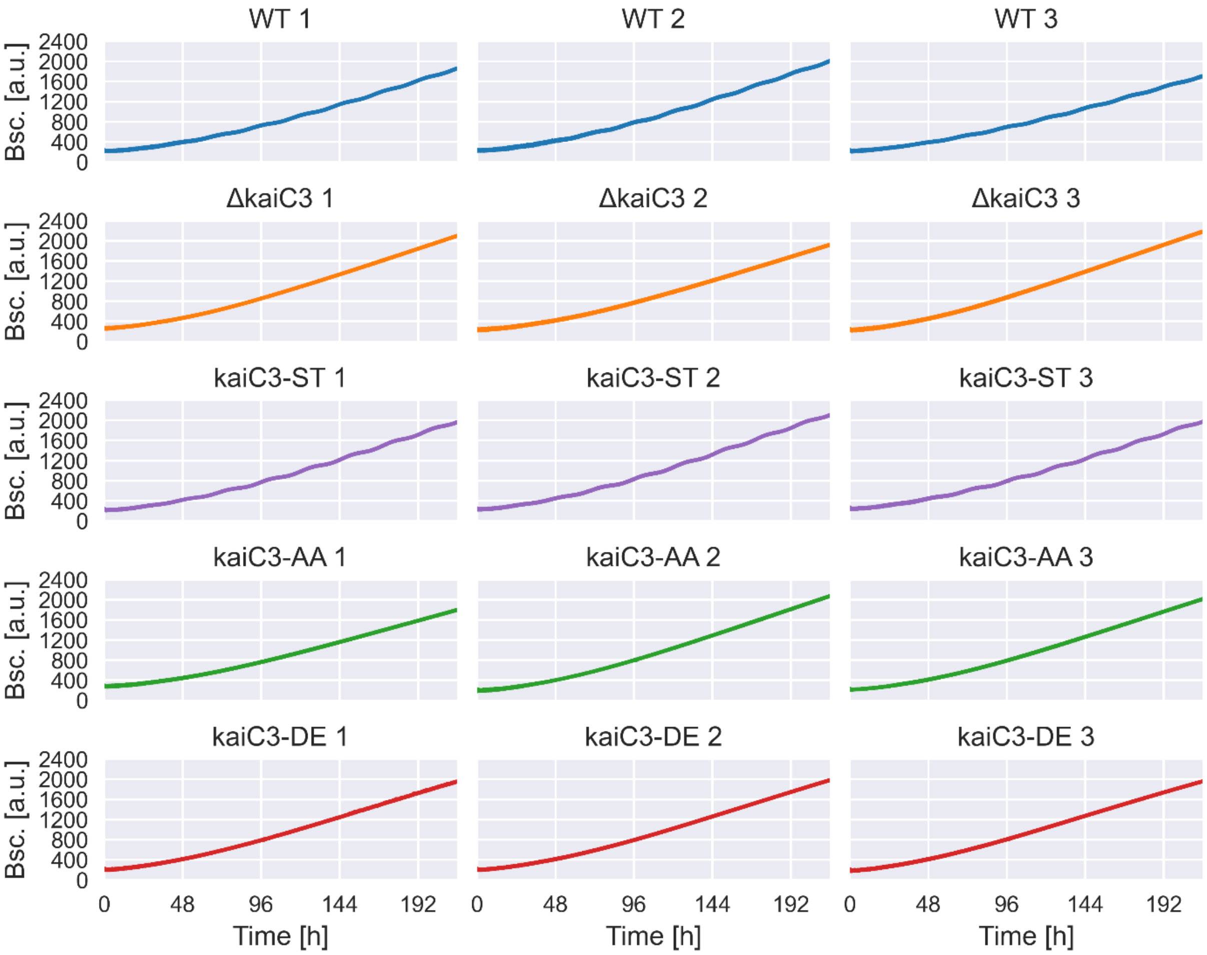


**Figure S9: Raw backscatter data for the individual replicates** of the strains WT (blue), Δ*kaiC3* (orange), *kaiC3*-ST (purple), *kaiC3*-AA (green), and *kaiC3*-DE (red) measured in batch cultures.

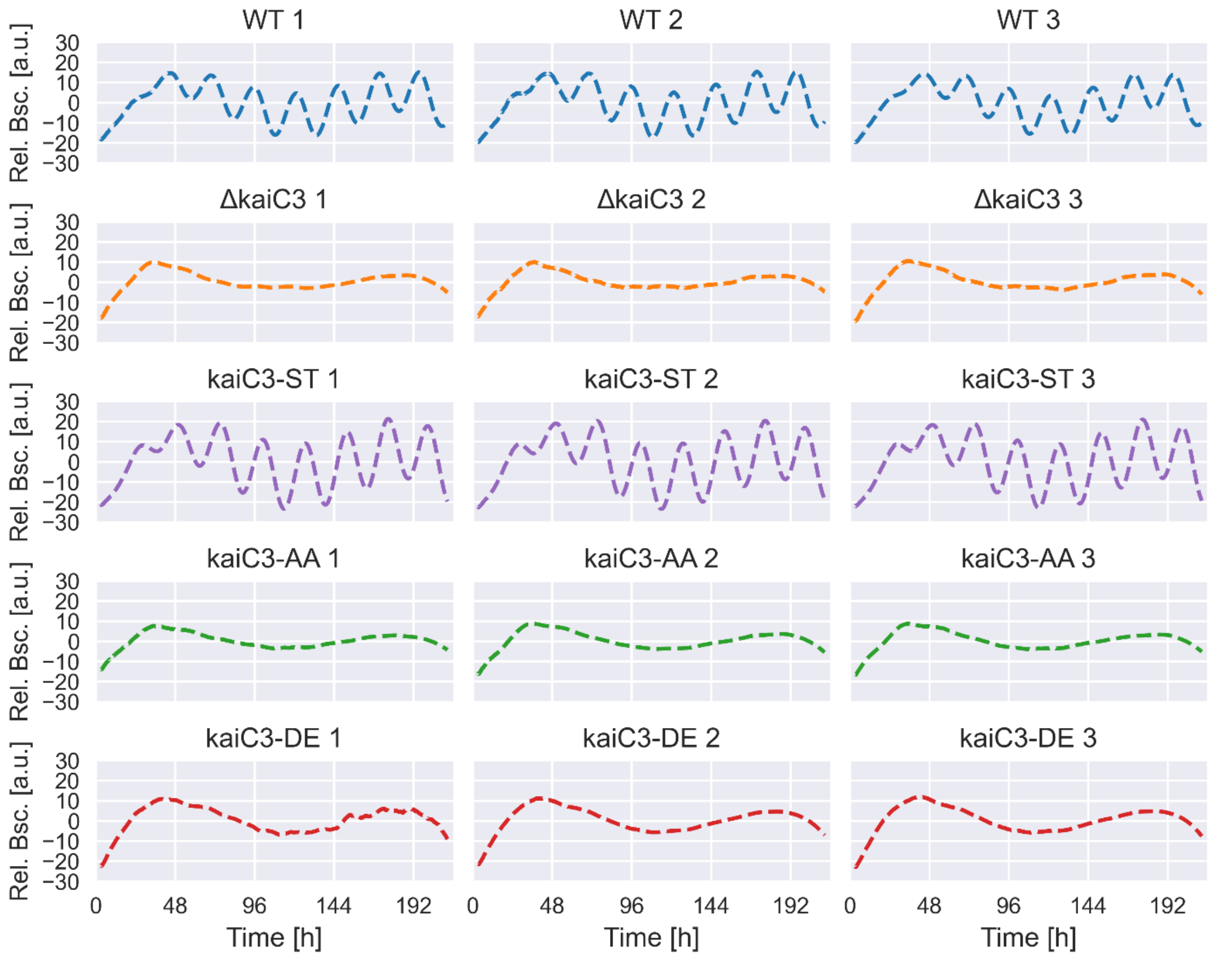


**Figure S10: Isolated smoothed backscatter oscillation for the individual** replicates of the strains WT (blue), Δ*kaiC3* (orange), *kaiC3*-ST (purple), *kaiC3*-AA (green), and *kaiC3*-DE (red) measured in batch cultures.

#### Strain generation

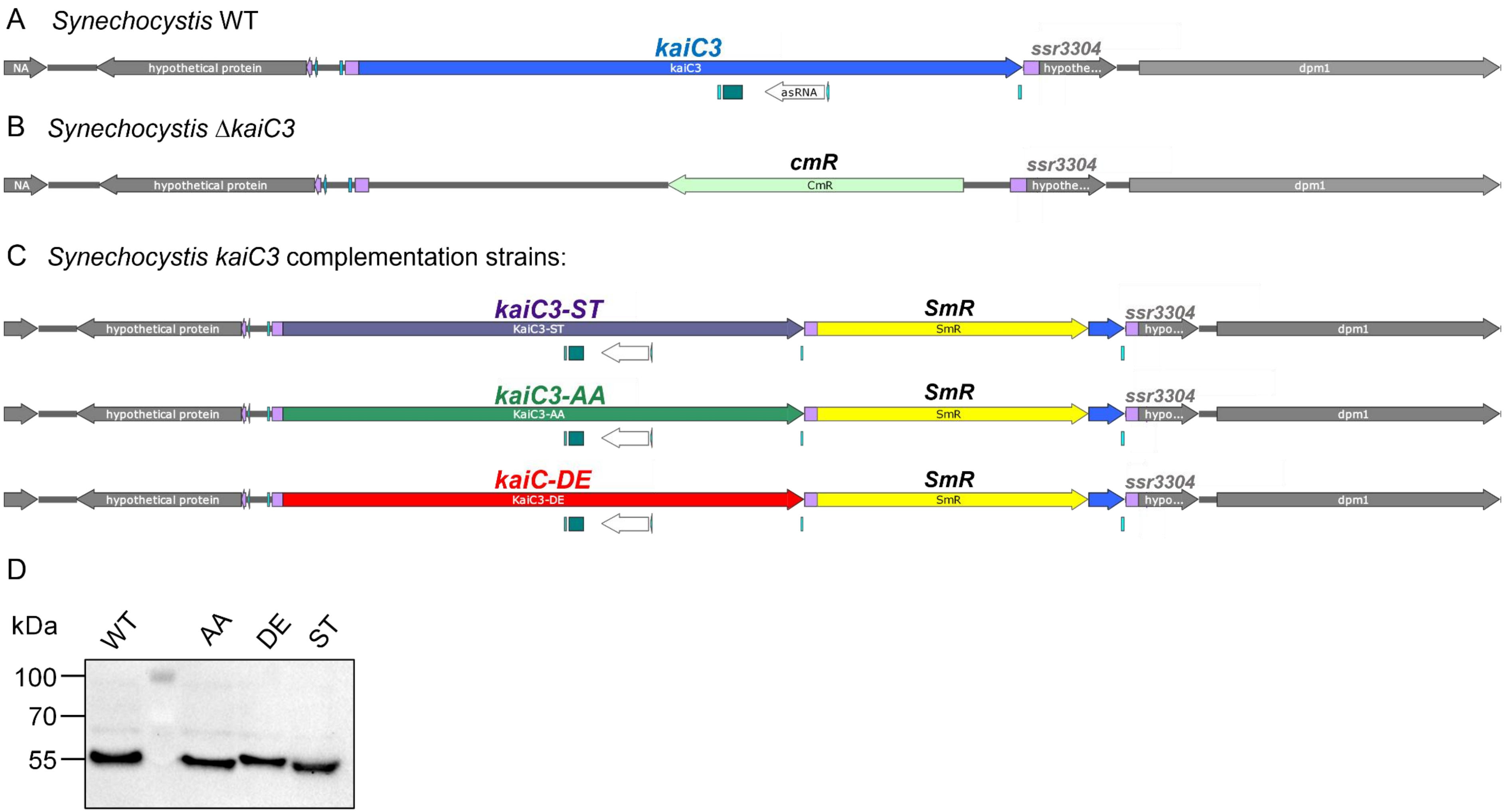


**Figure S11: Genomic context of *kaiC3* and generation of complementation strains.** A: According to the annotation of Mitschke et al., a TATA box (turquoise) is present at the end of the coding sequence (CDS) of *kaiC3* (blue) and the 5'UTR (violet) of the hypothetical protein Ssr3304 starts directly downstream of *kaiC3*, followed by the CDS of Ssr3304 (grey, 38 bp downstream of *kaiC3*) [1]. Accordingly, the region downstream of *kaiC3* may function as a terminator for *kaiC3* transcription, and the end of the *kaiC3* CDS may also function as a promoter/RBS for ssr3304. B: In *Synechocystis*

sp. PCC 6803 Δ*kaiC3* a CmR resistance cassette, together with a region of a plasmid, was inserted between the upstream and downstream flanking regions of *kaiC3* [2,3]. C: In the *kaiC3* complementation strains, the CmR cassette in *Synechocystis* sp. PCC6803 Δ*kaiC3* [3] was exchanged with the respective *kaiC3* coding sequences (*kaiC3*-ST, *kaiC3*-AA, or *kaiC3*-DE). To maintain the native terminator of *kaiC3*, we kept the 38bp downstream region of *kaiC3* and added a SpecR cassette as a selection marker behind it. We inserted the last 105 bp of *kaiC3* behind the SpecR cassette to maintain the hypothetical promoter of ssr3304 intact. Genomic maps were generated using the SnapGene software. D: *Synechocystis* sp. PCC 6803 WT and complementation strains (Δ*kaiC3* +*kaiC3*-AA, Δ*kaiC3* +*kaiC*3-DE, Δ*kaiC3*+*kaiC3*) were inoculated in 20 ml BG11 and grown at 150 rpm, 0.5 % CO2, and constant illumination with approximately 25 $\mu mol_{(photons)} \cdot m^{-2} \cdot s^{-1}$. On days 5 and 6, cells were diluted to an $OD_{750nm}$ of 0.4 and grown for two more days to $OD_{750nm}$ ~1.3-1.5. Cells from 4 ml culture were sheared with glass beads in lysis buffer (8M urea, 20mM HEPES pH 8.0). Whole cell extracts were normalized to 100 ng Chl a content, separated via SDS-PAGE using a 7.5% PAA gel, and subsequently blotted and immunodecorated with a KaiC3-specific antibody (1:3.750 in TBST) [4] and goat anti-rabbit IgG (H + L) Secondary Antibody, HRP (Thermo Fisher Scientific, 1:50,000, 1 h RT). After applying the Pierce SuperSignal West Pico detection reagent (Thermo Scientific), the signal was imaged using a ChemiDoc XRS+ Imaging System with ImageLab™ Software (BioRad).

**Table S3: List of primers used to generate the investigated strains.** The PaqCI recognition site (**cacctgc**) is indicated in bold. PaqCI cleaves the leading strand four bp downstream of the recognition site and the lagging strand eight bp downstream of the recognition site. In the table, the four bases following the recognition sequence are marked in italics. The complementary sticky ends produced after cleavage are marked in color, bold, and underlined. Primers were designed using the NEBridge Golden Gate assembly tool from New England biolabs.

| Primer | Sequence (5' → 3') | Purpose |
|---|---|---|
| 2423 | ggctac**cacctgc***tttc***aaag**gatgatcgaccaagagac | Amplification of *kaiC3*(-variants) from pASK-NStrep-*kaiC3*, *kaiC3*-AA or *kaiC3*-DE [5] (the sequence from start codon to Tyr434 is amplified) |
| 2424 | ggctac**cacctgc***attc***tcca**cgtaacgcaaaagaataatcgaatcggtaattgc | |
| 2425 | ggctac**cacctgc***attc***tgga**aatgtatggcgaaatg | Amplification of *kaiC3* (starting at Val435) and 41 bp downstream from genomic *Synechocystis* DNA |
| 2426 | ggctac**cacctgc***actt***ccgc**ttccctcatggattgattgcctccagg | |
| 2427 | ggctac**cacctgc***actt***gcgg**tgatcgccgaagtatc | Amplification of SpecR from pSHDY [6] |
| 2428 | ggctac**cacctgc***tgcc***acta**ccttggtgatctcgc | |
| 2429 | ggctac**cacctgc***tgcc***tagt**cggcaaataaggtaaacctttccgtaatg | Amplification of the last 105 bp of *kaiC3* plus 15bp downstream from genomic *Synechocystis* DNA |
| 2430 | ggctac**cacctgc***agga***ttac**aaagggaagaattgactatattttctcatcgaataaacc | |
| 2431 | ggctac**cacctgc***agga***gtaa**acctggaggcaatc | Amplification of backbone including flanking regions upstream of *kaiC3* and 15bp downstream of *kaiC3* from pJET-d*kaiC3*-CmR [2] |
| 2432 | ggctac**cacctgc***tttc***cttt**cactgccccatactc | |
